# A novel sweeping antibody exhibits efficient clearance of the cancer- and autoimmunity-associated cytokine interleukin 16

**DOI:** 10.1101/2025.07.29.666814

**Authors:** Jillian M. Baker, Alexander Wang, Kenneth A. Dietze, Djordje Atanackovic, Tim Luetkens

## Abstract

Targeting soluble antigens using conventional monoclonal antibodies is challenging due to high levels of antigen and limited antigen clearance per antibody molecule. Sweeping antibodies are engineered monoclonal antibodies that more efficiently clear soluble antigens than conventional antibodies. Sweeping antibodies contain two modifications: (1) pH-dependent antigen binding to facilitate lysosomal degradation of the targeted antigen while allowing antibody recycling and (2) enhanced neonatal Fc receptor (FcRn) engagement, resulting in 50-1,000-fold increased clearance of target antigen compared to conventional antibodies. The pleiotropic cytokine interleukin 16 (IL-16) has been proposed as a promising therapeutic target for monoclonal antibody therapy, due to its high expression and potential disease-promoting function in autoimmune diseases and cancer. Here, we develop the first fully human antibody as well as multiple sweeping antibodies targeting IL-16. We demonstrate that amenability to the introduction of pH-dependent binding into anti-IL-16 antibodies is correlated with epitope size and proximity to a positively charged IL-16 residue, informing future sweeping antibody development. We demonstrate that anti-IL-16 sweeping antibodies exhibit significantly increased antibody recycling and IL-16 degradation indicating that these molecules are a superior approach for the therapeutic targeting of IL-16 compared to conventional antibodies.

**GRAPHICAL ABSTRACT:** 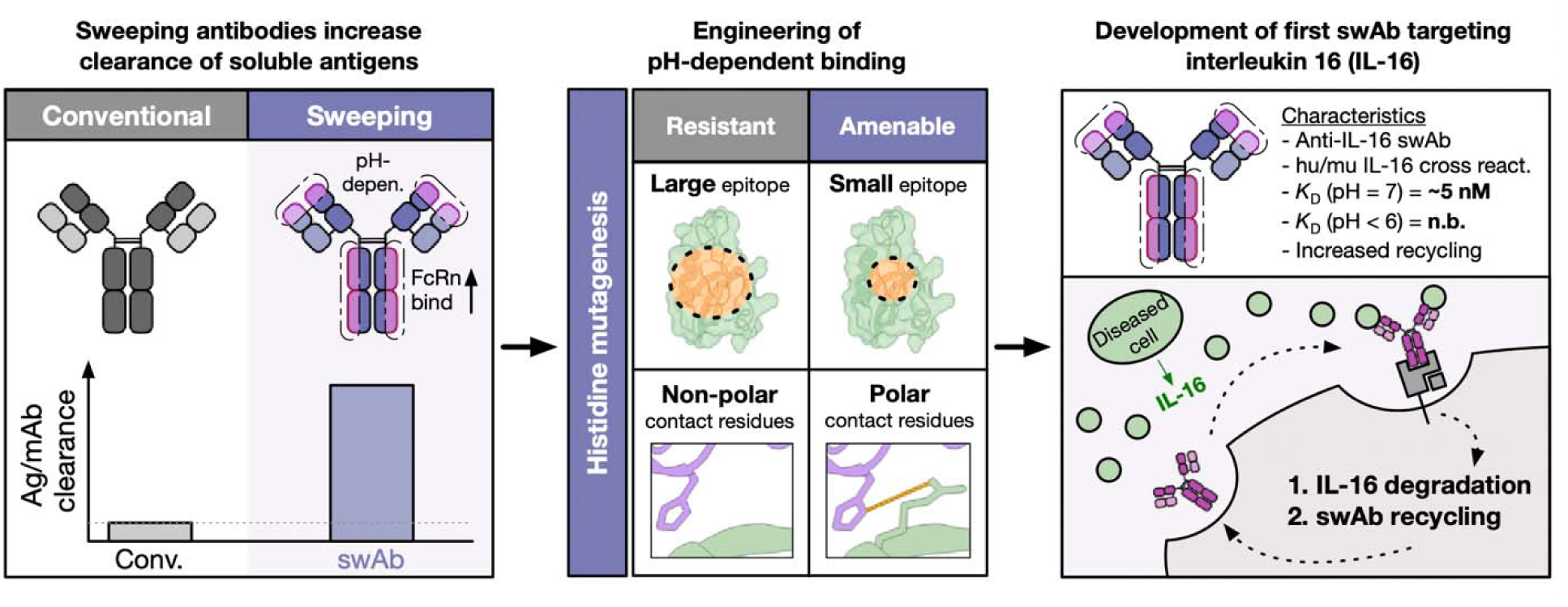

<u>One sentence summary:</u> Development of antibodies targeting IL-16 provides insights into sweeping antibody engineering and confers efficient clearance of the cancer- and autoimmunity-associated cytokine interleukin-16.

## INTRODUCTION

Monoclonal antibodies targeting cell surface antigens are an effective therapeutic approach across different clinical indications^1^. Similarly, targeting of some soluble ligands by antibodies has been shown to be feasible but targeting this class of proteins poses unique challenges^2^. Specifically, the high amount of antigen and its persistent production, i.e. in cancer and chronic diseases, requires high antibody levels and frequent dosing to achieve a therapeutic effect^3^. Sweeping antibodies have the potential to overcome these challenges by more efficiently clearing antigen, preventing antigen-antibody complex accumulation and thereby increasing the amount of antigen that can be cleared per antibody molecule. Sweeping antibodies differ from conventional antibodies in two ways: (1) sweeping antibodies exhibit pH-dependent binding to allow separation of the bound antigen in the sorting endosome and its subsequent selective degradation, and (2) they provide enhanced binding to the neonatal Fc receptor (FcRn) to facilitate more efficient antibody uptake and recycling^4^. Introducing these properties into antibodies adds a substantial layer of complexity from an engineering perspective but can result in antigen clearance 50-1,000-fold higher than wildtype (WT) antibodies^5^. The first sweeping antibody, ravulizumab^6^ targeting the complement protein C5 has recently been FDA-approved. Here, we develop novel sweeping antibodies targeting the antigen interleukin 16 (IL-16)^7,8^. IL-16 is a cytokine released by macrophages, eosinophils, mast cells, B cells and T cells following antigen, mitogen^9^, histamine^10,11^ or serotonin^12^ stimulation^13^. IL-16 has been shown to play a key role in the pathogenesis of allergy/asthma^14–16^, autoimmune diseases^17–19^ and cancer^20,21^. Therapeutic targeting of IL-16 may be an effective treatment for these diseases, however, its high serum levels may prove challenging to clear by conventional antibodies.

In this study, we generate sweeping antibodies against IL-16 by performing contact residue-targeted as well as scanning histidine mutagenesis of two anti-IL-16 antibodies. For one of the parental antibodies, this approach resulted in the successful elimination of antigen binding at lower pH and we show that the extent of pH dependency that can be introduced correlates with the size of the targeted epitope and proximity to a positively charged antigen contact residue. Importantly, we found that sweeping antibodies were able to degrade significantly higher amounts of IL-16 than the conventional parental antibody.

Our work provides 1) a roadmap to the identification of antibodies that are likely to be amenable to pH engineering and 2) the first sweeping antibodies against IL-16, a potential new therapeutic approach for the treatment of patients with autoimmune diseases and cancer.

## RESULTS

### Development and characterization of high-affinity anti-IL-16 antibodies

Bioactive interleukin-16 (IL-16) is a cytokine that is secreted by healthy lymphocytes (Fig. 1A) and different disease-associated cell types. It has previously been shown that IL-16 contributes to the pathogenicity of various malignant and autoimmune diseases (Fig. 1B). We hypothesize that sweeping antibodies are an effective approach for the neutralization and clearance of IL-16 for the treatment of human disease.

**Figure 1:**
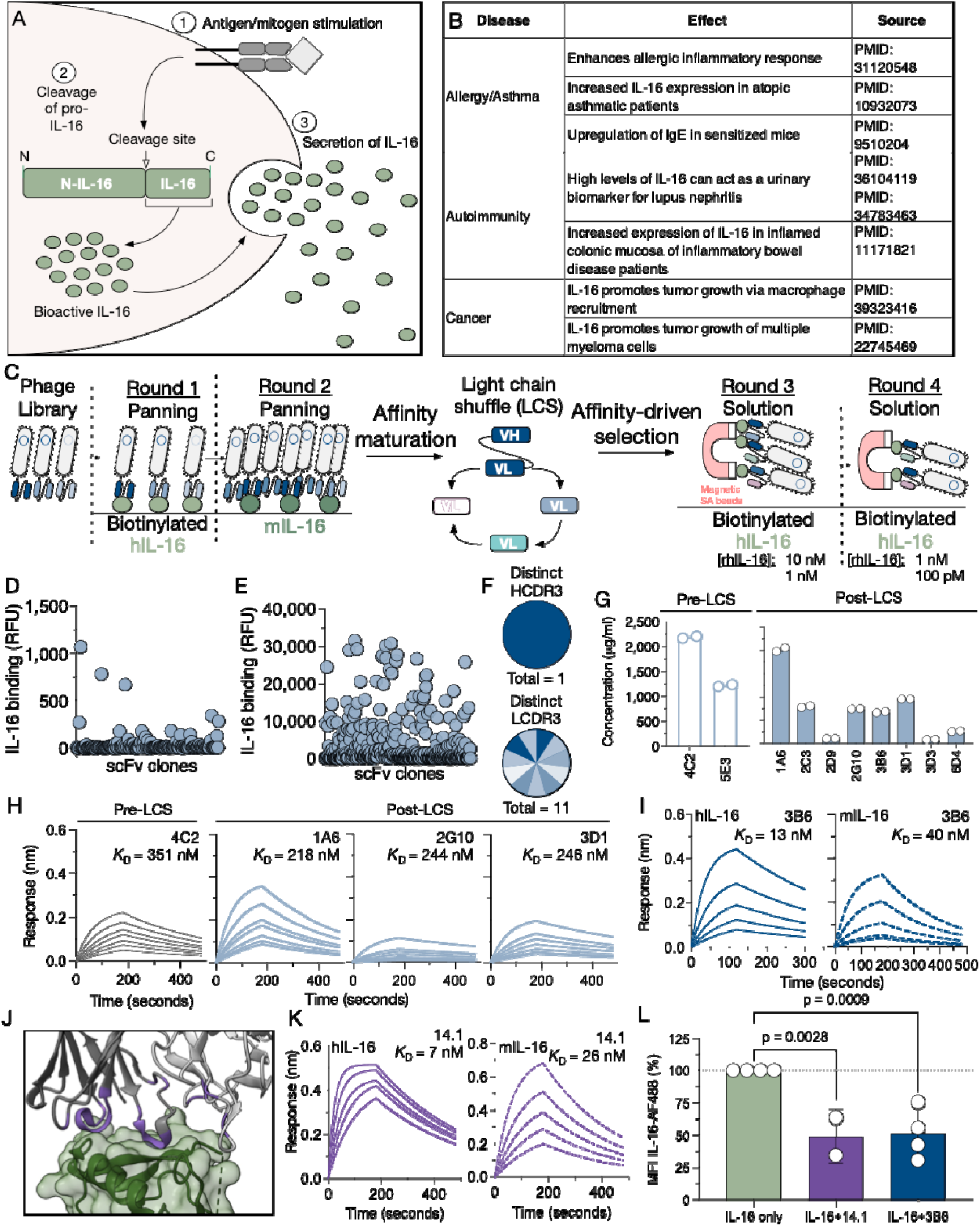
Development and characterization of high-affinity anti-IL-16 antibodies. **(A)** Schema of the intracellular processing and release of bioactive IL-16 from an IL-16-secreting cell. **(B)** Summary of the involvement of IL-16 in different human diseases. **(C)** Schema of the selection of a fully human anti-IL-16 antibody using single chain variable fragment (scFv) phage display. **(D)** Binding of round 2 (before light chain-shuffling) scFvs to human IL-16 (hIL-16) as determined by time-resolved fluorescence immunoassay (TRFIA). **(E)** Binding of round 3 and 4 (after light chain-shuffling and solution-phase selections) scFvs human IL-16 as determined by TRFIA. (**F**) Pie charts represent distinct heavy (HCDR3) and light (LCDR3) complementarity-determining region 3 sequences of single clones post-light-chain shuffling (LCS) as determined by Sanger sequencing. **(G)** Protein concentration of two pre-light chain shuffled (LCS) scFvs (white) and eight of the highest binding post-LCS (blue) scFvs after conversion into single-arm Fabs. Data indicate means ± SD from technical replicates (n = 2). **(H)** Bio-layer interferometry (BLI) sensorgrams of one pre-LCS Fab (black; 4C2) and three post-LCS Fabs (blue; 1A6, 2G10, 3D1) binding to hIL-16. Data indicate representative results from two independent experiments. Analyte concentrations used are summarized in Supplementary Table 1. **(I)** BLI sensorgrams of post-LCS Fab, clone 3B6, binding to human (hIL-16) and mouse IL-16 (mIL-16) at neutral pH. Data indicate representative results from two independent experiments. **(J)** Structure of the anti-IL-16 antibody 14.1 (grey, CDRs in purple) in complex with hIL-16 (green)^25^. **(K)** BLI sensorgrams of 14.1 Fab binding to hIL-16 and mIL-16 at neutral pH. Data indicate representative binding from two independent experiments. **(L)** Mean fluorescence intensity of CD3^+^ T cells stained with IL-16 conjugated to Alexa Fluor 488 (IL-16-AF488) only, IL-16-AF488 pre-incubated with 3B6 Fab, and IL-16-AF488 pre-incubated with 14.1 IgG. Data indicate means ± SD from individual experiments (n = 2-4). Data normalized to IL-16 only condition and reported as a percentage. Statistical differences between the experimental means and control mean (IL-16 only) were determined by ordinary one-way ANOVA followed by Dunnett’s multiple comparisons test.

To attempt development of sweeping antibodies targeting IL-16, we first generated a fully human anti-IL-16 antibody using antibody phage display (Fig. 1C). To enable phage selections, we expressed two variants of in vivo biotinylated human IL-16 (hIL-16) (Suppl. Fig. 1A). IL-16 with a flexible C-terminal linker had a significantly higher yield (Suppl. Fig. 1B) and was chosen for phage selections and all downstream assays. Sequential panning selections were performed on hIL-16 and commercially available mouse IL-16 (mIL-16). Binding of selected single-chain variable fragments (scFvs) was confirmed on a polyclonal (Suppl. Fig. 1C) and monoclonal level (Fig. 1D). To enhance binding to hIL-16, light-chain shuffling^22,23^ followed by solution-phase selection with limiting amounts of IL-16 was performed. Monoclonal scFvs demonstrated binding to IL-16 in a time-resolved fluorescence immunoassay (TRFIA) (Fig. 1E). Post-light chain-shuffled scFvs displayed 10-50x higher binding to IL-16 compared to pre-light chain-shuffled scFvs (Suppl. Fig. 1D). All high binding clones shared the same heavy chain CDR3 but had unique light chain CDR3s (Fig. 1F). Ten of the eleven clones and two pre-light chain shuffled clones were converted to and expressed as single-arm Fabs (Fig. 1G, Suppl. Fig. 1E). Using bio-layer interferometry (BLI), we found that 5 clones (4C2, 1A6, 2G10, 3D1 and 3B6) retained binding to hIL-16 once converted to Fab format (Fig. 1H/I). One clone, 3B6, had low nanomolar affinity to hIL-16 (K*_D_* = 13 nM) and showed cross-reactivity (Suppl. Fig. 1F) to mIL-16 (K*_D_*= 40 nM) (Fig. 1I) but not to an irrelevant control protein (Suppl. Fig. 1G). In addition to 3B6, for our subsequent work, we also included a previously identified IL-16 mouse monoclonal antibody, clone 14.1^8^. 14.1 binds to soluble IL-16 (Fig. 1J) and has previously been shown to have some therapeutic activity in an animal model of acute kidney injury^24^. We show that, similar to 3B6, 14.1 has low nanomolar affinity to human (K*_D_* = 7 nM) and mouse IL-16 (K*_D_* = 26 nM) (Fig. 1K, Supp Fig. 1H) and both antibodies efficiently prevent binding of fluorescently labeled IL-16 to activated T cells (Fig. 1L). Because of their favorable biophysical and functional properties, both antibody clones were included for downstream sweeping antibody engineering.

### Parental anti-IL-16 antibodies show minimal pH-dependent binding

The predominant mechanism-of-action of conventional therapeutic antibodies against soluble antigens is the binding of target antigen with high affinity to block interaction of the target with its receptor^26^. In principle, the capacity of an antibody to neutralize target antigen could be amplified if, instead of blocking antigen-receptor binding, the antibodies preferentially engaged in antigen degradation followed by antibody recycling. However, conventional antibodies are stoichiometrically restricted in this context because they are cleared together with the bound antigen^27,28^. Sweeping antibodies circumvent this limitation of conventional antibodies by avoiding lysosomal degradation and allowing efficient antibody recycling. This is achieved through the introduction of two features that distinguish sweeping antibodies from conventional antibodies: (1) increased affinity of the Fc domain to FcRn, enhancing internalization at neutral pH^5^ and antibody recycling, and (2) pH-dependent binding to antigen (Fig. 2A). Following internalization, in the acidic environment of the sorting endosome, sweeping antibodies dissociate from the antigen and are recycled back out to the extracellular space with unoccupied binding domains to allow binding of new antigen molecules, substantially increasing their clearance capacity over conventional antibodies^29^.

**Figure 2:**
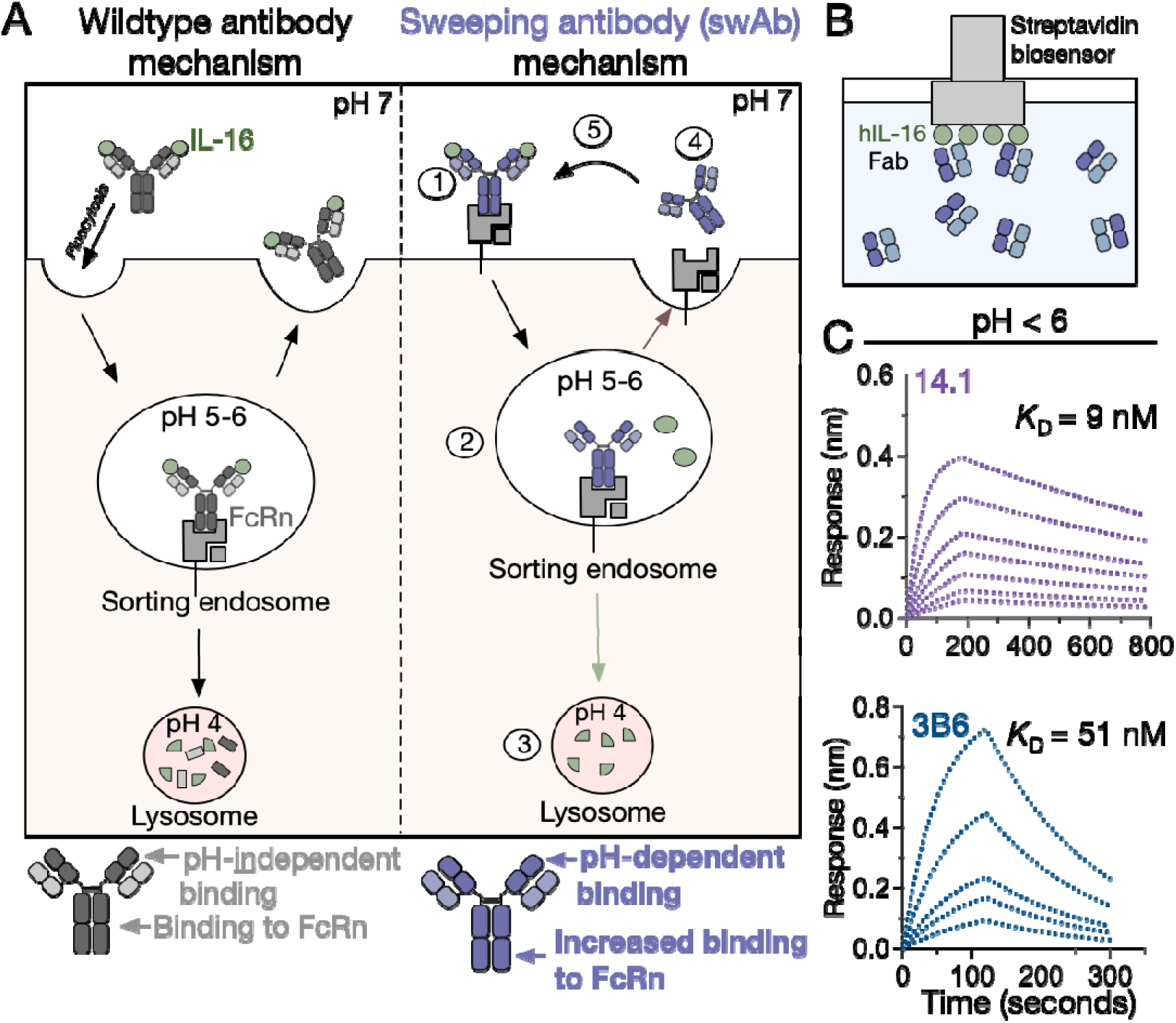
pH-dependency of parental anti-IL-16 antibodies. **(A)** Schema of differences in the internalization and recycling of conventional and sweeping antibodies in neonatal Fc-receptor (FcRn)-expressing cells. **(B)** Schema of the bio-layer interferometry (BLI) assay set-up for measuring Fab (analyte) binding to hIL-16 (ligand) at neutral and acidic pH (pH < 6). **(C)** BLI sensorgrams of 14.1 Fab and 3B6 Fab binding to hIL-16 at acidic pH. Data indicate representative binding from two independent experiments.

To engineer anti-IL-16 sweeping antibodies, we first determined the degree of pH-dependent binding to IL-16 by the two parental anti-IL-16 antibodies. BLI was used to determine the affinity of the antibodies to hIL-16 at neutral pH and at pH < 6 in two assay orientations (Fig. 2B, Suppl. Fig. 2A). The orientation in which hIL-16 served as ligand and the single-arm Fab as analyte was chosen for downstream analyses (Fig. 2B, Suppl. Fig. 2B). Both 14.1 and 3B6 show high affinity for hIL-16 at neutral pH (14.1: *K*_D_ = 7 nM; 3B6: *K*_D_ = 13 nM) and at pH < 6 (14.1: *K*_D_ = 9 nM; 3B6: *K*_D_ = 51 nM, Fig. 2C), demonstrating relatively pH-independent binding by the parental clones.

### Generation of pH-dependent anti-IL-16 antibodies

The histidine side chain has a pKa of ∼6.6^30^. When located in critical binding positions, histidine residues can render a protein an acid switch. Acid switches allow antigen binding in the deprotonated state (pH > 7) while preventing binding in the protonated state (pH < 6)^5,31^. To introduce pH-dependent binding, we used different approaches for each of the parental antibodies. For 3B6, each amino acid residue in the complementarity-determining regions (CDRs) in the heavy and light chains was replaced with histidine, creating single-histidine heavy and light chain variants (1xH). For 14.1, to generate pH-dependent clones, we replaced the previously described^25^ contact residues in the heavy chain and light chain with histidine residues creating single-histidine variants (1xH). 3B6 1xH Fabs and 14.1 1xH IgGs were expressed in mammalian cells and binding to IL-16 was assessed at neutral pH (Fig. 3A/B, Suppl. Fig 3A-C). Compared to the parental clone, most 14.1 1xH variants retained expression except for variants with histidine residues in the heavy chain CDR3, which showed markedly reduced expression. A similar pattern was also observed in the 3B6 variants with mostly those clones containing histidine replacements in the heavy and light chain CDR3s showing reduced expression. Almost half of 14.1 (8/18) and 3B6 (23/49) 1xH variants showed substantially reduced binding to hIL-16 at neutral pH compared to the respective parental clone (Fig. 3A/B). The other single-histidine variants showed similar or increased binding to hIL-16 compared to the respective parental clone (Fig. 3A/B).

**Figure 3:**
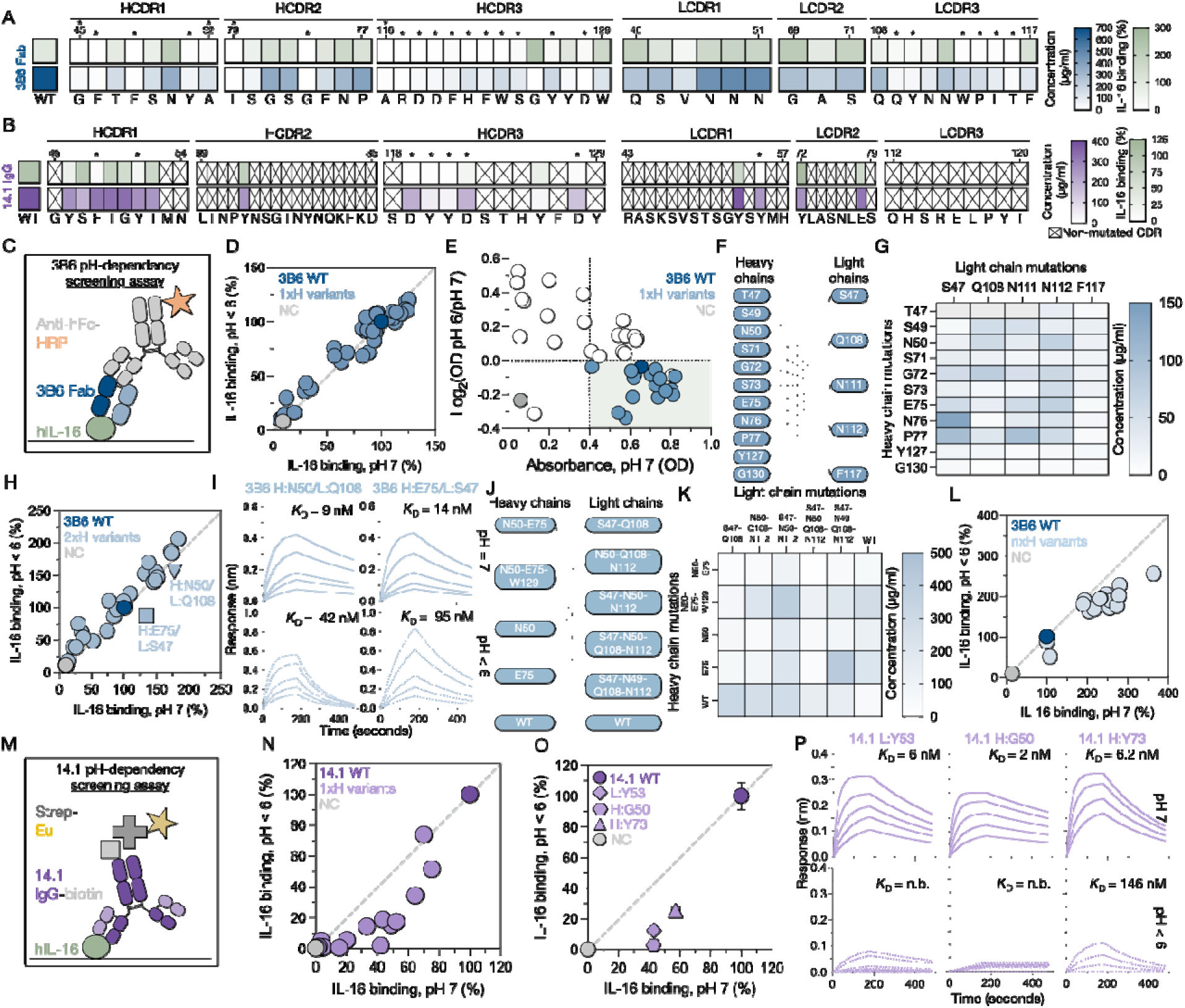
Generation of pH-dependent antibodies from parental high-affinity antibodies using histidine mutagenesis. **(A)** 3B6 and **(B)** 14.1 single histidine variant Fab protein concentrations and hIL-16 binding at neutral pH as a percentage of parental antibody binding as determined by ELISA. The residues in parental heavy (HCDR) and light (LCDR) chain complementarity-determining regions that were mutated to histidine are indicated below. Asterisks indicate variants with < 25% binding compared to parental 3B6. Heatmap data represent averages from technical replicates (n = 2-3). WT = wildtype/parental. Crossed out boxes indicate non-contact residues that were not mutated. **(C)** Schema of ELISA setup used to determine pH-dependency of 3B6 Fab histidine variants. **(D)** Binding of single-histidine 3B6 variants (1xH) to hIL-16 at neutral pH (pH 7) and acidic pH (pH < 6). Data are normalized to parental 3B6 (WT) binding at both pH values and reported as a percentage; negative control (NC) = 4% BSA/PBS. Dotted line represents relative pH-dependency comparable to parental 3B6. **(E)** Difference in relative binding at acidic/neutral pH vs. absolute binding to hIL-16 at neutral pH by 3B6 1xH variants as determined by ELISA. **(F)** Schema representing the pairing of variant heavy chains with variant light chains for double histidine variants (2xH). The notation indicates amino acid and position in parental (WT) 3B6 that was mutated to histidine. **(G)** Heatmap of protein concentrations of 3B6 2xH Fab variants. Data represents average of technical replicates (n = 2). **(H)** Binding of 3B6 2xH variants to hIL-16 at neutral and acidic pH. Data normalized to parental 3B6 (WT) binding at the respective pH; negative control (NC) = 4% BSA/PBS. Dotted line represents relative pH-dependency comparable to parental 3B6. **(I)** Bio-layer interferometry (BLI) sensorgrams of 3B6 H:N50/L:Q108 and H:E75/L:S47 binding to hIL-16 at neutral and acidic pH. **(J)** Schema representing the pairing of combinatorial variant heavy chains with combinatorial variant light chains for multi-histidine variants (nxH). **(K)** Heatmap of protein concentrations of nxH 3B6 Fab variants. Data represents average of technical replicates (n = 2). **(L)** Binding of nxH 3B6 variants to hIL-16 at neutral and acidic pH as determined by ELISA. Data normalized to parental 3B6 (WT) binding at both pH values; NC = 4% BSA/PBS. Dotted line represents relative pH-dependency comparable to parental 3B6. **(M)** Schema of TRFIA setup used to determine pH-dependency of 14.1 IgG histidine variants. **(N)** Screening and **(O)** validation assays to determine binding of single-histidine 14.1 variants (1xH) to hIL-16 at neutral and acidic pH, as determined by TRFIA. Data normalized to parental 14.1 (WT) binding at the respective pH; NC = 4% BSA/PBS. Dotted line represents relative pH-dependency comparable to parental 14.1. **(O)** Data represents average of technical replicates (n = 3). **(P)** BLI sensorgrams of selected 14.1 1xH variants binding to hIL-16 at neutral and acidic pH. Data indicate representative binding from two independent experiments.

We next determined whether introduction of histidine residues increased the respective clones’ pH-dependent binding. To determine pH-dependency of 3B6 1xH variants, binding assays were performed at both neutral pH and at pH < 6 (Fig. 3C). None of the 3B6 1xH variants displayed distinct pH-dependent binding to hIL-16 compared to parental 3B6 (Fig. 3D). To enhance pH-dependency, 1xH mutations that retained binding to hIL-16 at neutral pH (OD_pH 7_ > 0.4) and introduced at least some pH-dependency (OD_pH 6_/OD_pH 7_) < 1) were selected for the generation of combinatorial variants (Fig. 3E). Every selected 1xH heavy chain variant was paired with most selected 1xH light chain variants to create 2xH variants (Fig. 3F). 2xH variants were expressed as single-arm Fabs (Fig. 3G), isolated, and used in binding assays. Similar to the 1xH variants, the 2xH variants also did not display substantially increased pH-dependent binding to hIL-16 (Fig. 3H). Two variants (H:N50/L:Q108 and H:E75/L:S47) showed the most pH-dependent binding in the 2xH screen, which was confirmed using BLI (Fig. 3I). However, the 2xH screening still only resulted in a candidate with a ∼7-fold difference in affinity between neutral pH and pH < 6. This prompted us to combine multiple histidine mutations on each chain to potentially achieve a greater reduction in binding at low pH. Each new heavy chain was paired with new light chains to create nxH variants (Fig. 3J). In addition to choosing histidine mutations with the highest pH-dependency, we also selected residues that represented sequence liabilities to enhance the biophysical properties of the resulting antibodies^26^. All nxH variants were expressed as single-arm Fabs (Fig. 3K, Suppl. Fig. 3D). Overall, these clones showed improved expression compared to the 2xH mutants likely due to the removal of sequence liabilities^32^. Next, we determined pH-dependency of nxH variants. Strikingly, almost all nxH variants showed enhanced binding to hIL16 at neutral pH and pH < 6 (Fig. 3L) and all were found to have low nanomolar *K*_D_ at both pH levels (Suppl. Fig. 3E). We concluded that, despite achieving some degree of pH-dependent binding with two histidine mutations, introducing additional histidine residues did not yield increased pH dependency.

We next explored whether 14.1 would be more amenable to the introduction of pH dependency. We first performed binding assays of 1xH variants at neutral pH and pH < 6 (Fig. 3M). Of the 8 variants that retained binding at neutral pH, 7 now showed pH-dependent binding in our screening assay (Fig. 3N). We selected the three clones with the highest degree of pH dependency, one light chain variant (L:Y53) and two heavy chain variants (H:Y73,H:G50). In our validation assay, we confirmed retained binding of the three candidate antibodies to hIL-16 at neutral pH and reduced binding at pH < 6 (Fig. 3O). The three candidate antibodies were converted to single-arm Fabs to determine binding kinetics using BLI. Compared to parental 14.1, all three variants had retained similar binding kinetics at neutral pH (L:Y53: *K*_D_ = 6 nM, H:Y73: *K*_D_ = 6.2 nM, H:G50: *K*_D_ = 2 nM) (Fig. 3P). However, at pH < 6, binding was not detectable anymore by BLI for two variants (L:Y53, H:G50) and variant H:Y73 had a 24x reduction in affinity (*K*_D_ = 146 nM).

Taken together, 3B6 appeared relatively resistant to the introduction of pH-dependency but in clone 14.1, replacement of single amino acids with histidine led to multiple variants with substantial pH-dependent binding. We hypothesize that overall differences in the antibody/antigen interface may render some antibodies amenable and others resistant to the introduction of pH-dependent binding.

### Differences in target epitopes correlate with amenability to introduction of pH sensitivity

To better understand why 14.1 but not 3B6 was amenable to the introduction of pH dependency, we further characterized the interaction between the two antibodies with hIL-16. Using tandem BLI, we found that there was some competition between the clones (Fig. 4A), indicating that the epitopes for 3B6 and 14.1 are distinct but partially overlap.

**Figure 4:**
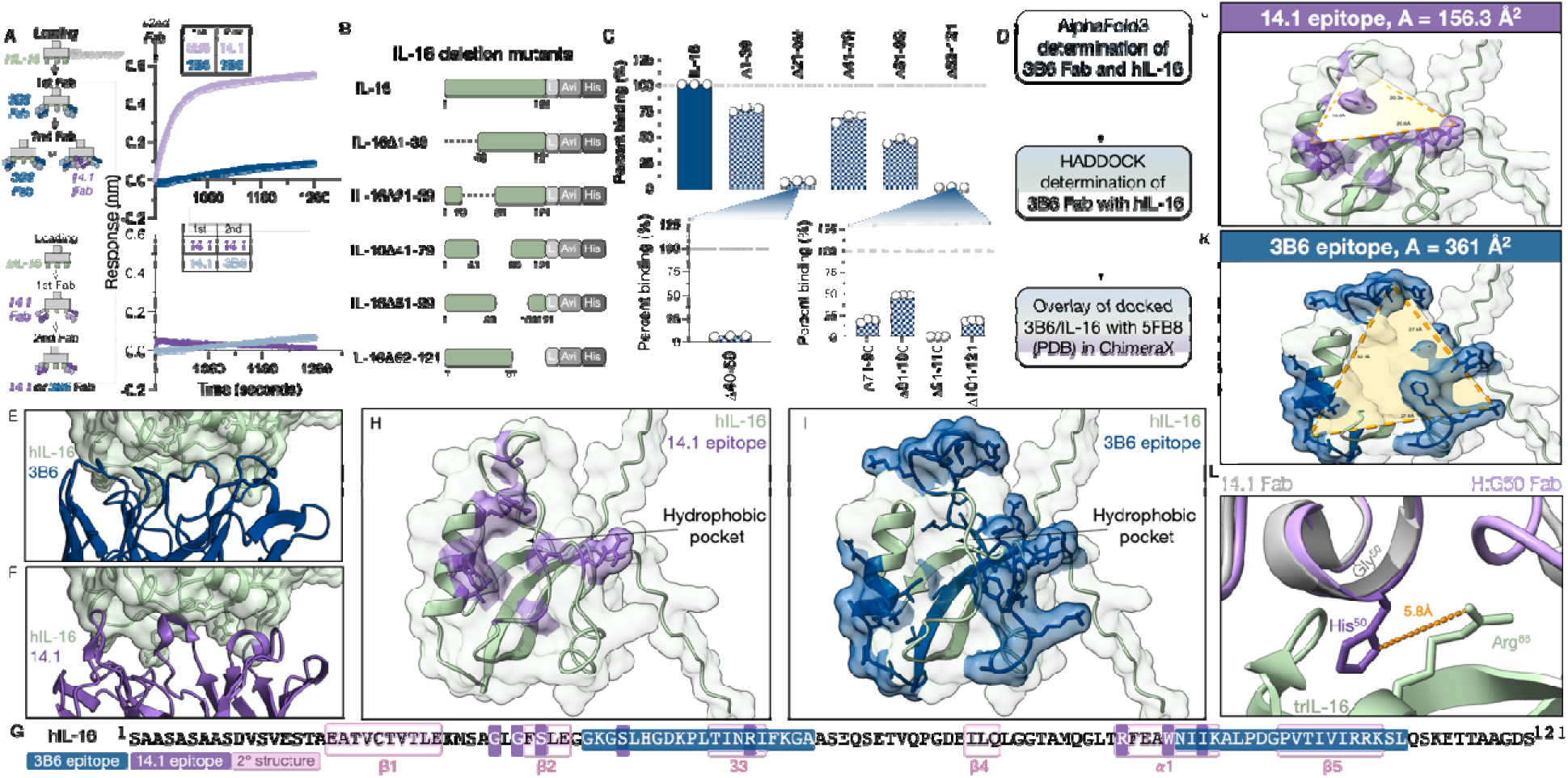
Differences in target epitopes confer amenability to introduction of pH sensitivity. **(A)** Binding of 3B6 or 14.1 to hIL-16 previously loaded with 3B6 or 14.1 Fabs as determined by tandem BLI. Data indicate representative binding from three independent experiments. **(B)** Schema of 40-amino acid, overlapping hiL-16 deletion mutants. **(C)** Binding of 3B6 IgG to wildtype hIL-16 or hIL-16 deletion mutants. Data indicate means ± SD from technical replicates (n = 3). **(D)** Schema of workflow to generate 3B6/hIL-16 complex structure for comparison to 14.1/hIL-16 complex. Structures of the interfaces between hIL-16 (green) and **(E)** 3B6 Fab (blue) or **(F)** 14.1 Fab (purple). **(G)** Sequence of hIL-16 with mapped epitope residues for 3B6 (blue), 14.1 (purple) and secondary structures (pink). hIL-16 (green) epitope residues for **(H)** 14.1 (purple) or **(I)** 3B6 (blue). Triangular area of the **(J)** 14.1 epitope or the **(K)** 3B6 epitope. **(L)** Structure alignment of 14.1 H:G50 (purple) with PDB: 5FB8 containing 14.1 (grey) and truncated hIL-16 (trIL-16) (green) showing the distance between the introduced histidine of H:G50 and Arg^86^ of trIL-16.

The epitope of 14.1 has previously been determined using X-ray crystallography^25^. To experimentally determine the epitope of 3B6, we first performed linear peptide scanning using overlapping 15-mer peptides (Suppl. Fig 4A). No significant binding of 3B6 to the linear peptides was observed indicating that the epitope of 3B6 may be conformational. We next expressed and purified 5 hIL-16 constructs with 40-amino acid overlapping deletions (Fig. 4B, Suppl. Fig 4B/C). Two of these deletion mutants displayed substantially reduced binding to IL-16 (Fig. 4C). To further narrow down the 3B6 epitope, we generated five 20-amino acid hIL-16 deletion mutants and found that only two of these mutants showed reduced binding: Δ40-59 and Δ91-110, corresponding to the β3-sheet and the α1-helix/β5-sheet of IL-16 (Fig. 4C, Suppl. Fig 4D/E), which was consistent with a conformational epitope. Using this information, we generated an in silico model of the 3B6:hIL-16 complex (Fig. 4D). We found that the overall orientation of 3B6 bound to hIL-16 (Fig. 4E) mirrored that of 14.1 (Fig. 4F), which is in line with our observation of a partially overlapping epitope. This was further confirmed by the mapping of the experimentally determined epitope residues onto the hIL-16 sequence (Fig. 4G).

We next explored potential reasons for the difference in our ability to introduce pH sensitivity. We first hypothesized that differences in the overall electrostatic potential of the antibodies’ respective epitopes could lead to increased repulsive charges differentially affecting binding at low pH. We therefore assessed the electrostatic potential of the epitopes of 14.1 and 3B6 at pH 7 and pH 5 (Suppl. Fig. 5). Both epitopes contained residues exhibiting strong positive electrostatic potential at pH 7 with basicity for residues slightly increasing at pH 5, without clear differences between the epitopes in terms of overall electrostatic potential.

We next considered if the size of the targeted epitopes could affect resistance to pH engineering using histidine mutagenesis. Specifically, we hypothesized that a larger epitope surface could be more resistant to this engineering approach due to a higher number of contact residues providing redundancy in antigen binding. Comparing the epitope areas of the two clones, we found that 14.1 has a narrow epitope with residues forming a tight hydrophobic pocket around the protruding heavy CDR3 loop (Fig. 4H). In contrast, the epitope area of 3B6 is more extensive and surrounds the same hydrophobic pocket but at a greater distance from its center (Fig. 4I). We found that the total area of the 3B6 epitope was more than twice as large as that of 14.1 (Fig. 4J/K). These data indicate that total epitope area may be an key parameter in conferring resistance to the introduction of pH sensitivity.

For 14.1, we had observed a stark increase in pH sensitivity following the histidine substitution of a single residue, H:G50. To better understand how this mutation confers this change in binding at low pH, we analyzed the structure of the complex between 14.1-H:G50 and hIL-16. At the site of substitution, the backbones of 14.1 and 14.1-H:G50 are predicted to overlap. However, replacement of Gly^50^ in 14.1 with histidine results in the introduction of a bulkier, positively charged residue at the interface between Fab and hIL-16 (Fig. 5L), only 5.8 Å from the positively charged hIL-16 contact residue, Arg^86^. It is likely that the H:G50 substitution confers electrostatic disruption of 14.1:hIL-16 binding at low pH.

**Figure 5.**
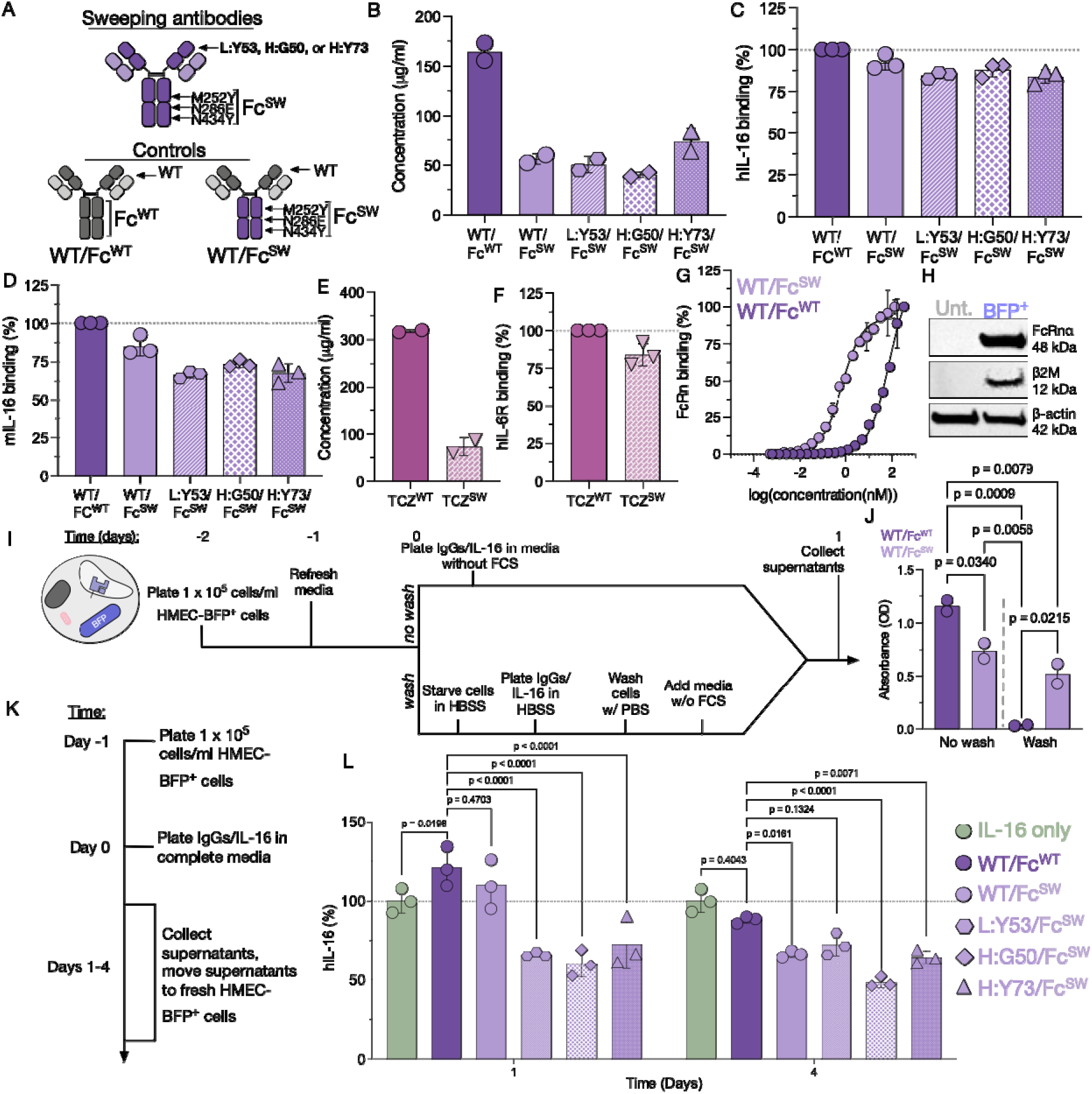
Anti-IL-16 sweeping antibodies efficiently target IL-16 for degradation. **(A)** Schema showing anti-IL-16 sweeping antibodies containing histidine substitutions as well as Fc mutations to enhance FcRn binding as well as pH-independent control antibodies with or without Fc mutations to enhance FcRn binding. **(B)** Protein concentrations of control and sweeping 14.1 antibodies. Data indicate means ± SD from technical replicates (n = 2). Binding of sweeping and control anti-IL-16 antibodies to **(C)** human and **(D)** mouse IL-16 as determined by ELISA. Data indicate means ± SD from technical replicates (n = 3). **(E)** Protein concentration of TCZ^WT^ and TCZ^SW^ antibodies. Data indicate means ± SD from technical replicates (n = 2). **(F)** Binding of TCZ^WT^ and TCZ^SW^ antibodiesto human IL-6R (hIL-6R) as determined by ELISA. Data indicate mean ± SD from technical replicates (n = 3). **(G)** Binding of WT/Fc^WT^ and WT/Fc^SW^ 14.1 antibodies to biotinylated human FcRn as determined by ELISA. Data indicate mean ± SD from technical replicates (n = 2). **(H)** Protein expression of FcRnα, β2 microglobulin and β-actin (housekeeping protein) in untransduced (unt.) and transduced HMEC cells. Data are representative of two independent experiments. **(I)** Schema of experimental setup to determine antibody recycling by HMEC-FcRn-β2M cells. **(J)** Sweeping antibody concentrations in the supernatant of HMEC-FcRn-β2M before and after washing. Data indicate means ± SD from technical replicates (n = 2). Statistical differences between the means were determined using an ordinary one-way ANOVA followed by Tukey’s multiple comparisons test. **(K)** Schema of experimental setup to determine IL-16 degradation in the presence of 14.1 control and sweeping antibodies and HMEC-FcRn-β2M cells. **(L)** hIL-16 levels after incubation with 14.1 control and sweeping antibodies and HMEC-FcRn-β2M cells for one or four days, as determined by ELISA. Data indicate mean ± SD from technical replicates (n = 3). Data are normalized to IL-16 only condition. Statistical differences were determined by two-way ANOVA followed by Šídák’s multiple comparisons test.

Taken together, our data show that 3B6 and 14.1 bind the same general region of IL-16 resulting in a partial epitope overlap and that their epitopes do not exhibit major differences in overall electrostatic potential. However, our data indicate that their difference in epitope area as well as favorable positioning of the 14.1-H:G50 residue in close proximity to a positively charged contact residue in the hIL-16 epitope may be key determinants in their amenability to the introduction of pH sensitivity.

### Anti-IL-16 sweeping antibodies efficiently target IL-16 for degradation

Having identified multiple 14.1-based pH-dependent binders, we next generated full sweeping anti-IL-16 antibodies by introducing three mutations (M252Y, N286E, and N434Y) to the Fc domain (Fc^SW^) to enhance binding to the neonatal Fc receptor (FcRn) (Fig. 5A)^5^. Increased binding of sweeping IgGs to FcRn has been shown to improve antibody internalization and subsequent clearance of antigen^4^. We generated wildtype 14.1 (parental/WT variable region + Fc^WT^), sweeping IgGs (pH-dependent variable region + Fc^SW^), and a control construct which contained the parental variable region and the FcRn-binding mutations (WT/ Fc^SW^). We found that introduction of the Fc domain mutations substantially reduced expression of the sweeping constructs compared to parental 14.1 (Fig. 5B, Suppl. Fig. 6A) but did not alter binding to hIL-16 (Fig. 5C) and mIL-16 (Fig. 5D). To determine whether the reduced expression of full sweeping antibodies was due to the sweeping Fc mutations and not a combination of sweeping Fc mutations and anti-IL-16 variable regions, we expressed a pair of established parental (TCZ^WT^) and sweeping (TCZ^SW^) anti-IL-6R antibodies targeting soluble IL-6R, based on the FDA-approved anti-IL-6R antibody, tocilizumab^5,33^. Similarly to 14.1 sweeping antibodies, TCZ^SW^ showed reduced expression compared to TCZ^WT^ (Fig. 5E) while retaining binding to IL-6R (Fig. 5F). These data confirm that the introduced Fc mutations reduce protein expression independent of target antigen. We next determined the effect of Fc mutations on binding to recombinant FcRn. We found that the introduced Fc mutations substantially increased binding to FcRn compared to the wildtype Fc region (Fig. 5G).

Finally, to evaluate overall sweeping antibody functionality, we used a combination of previously established recycling assays^34^ and newly designed degradation assays. For these assays, we engineered human microvascular endothelial cells (HMEC) using lentiviral transduction (Suppl. Fig. 6C) to overexpress FcRn alpha (FcRnα) and beta-2-microglobulin (β2M) together with a blue fluorescent protein (BFP) reporter (Suppl. Fig. 6D). Transduced HMEC-FcRnα-β2M cells were sorted based on BFP expression and expression of FcRnα and β2M in the sorted cells was confirmed by western blot (Fig. 5H). To determine how FcRn-binding enhancing Fc mutations affect internalization of the antibodies by HMEC-FcRnα-β2M cells, we used a modified version of a previously established recycling assay^34^ (Fig. 5I). In brief, HMEC-FcRnα-β2M cells were incubated with parental and sweeping antibodies and supernatants were harvested before and after washing. We observed reduced amounts of sweeping anti-IL-16 antibodies in supernatants from non-washed cells, indicating increased internalization. Following washes, we did not detect any parental antibodies but high levels of sweeping antibodies (Fig. 5J). These data are consistent with increased internalization by sweeping antibodies followed by efficient recycling whereas wildtype antibodies showed limited cell binding and no recycling. We next set out to determine the ability of anti-IL-16 sweeping antibodies to degrade IL-16. Parental and sweeping anti-IL-16 antibodies were incubated with IL-16 and HMEC-FcRn-β2M cells for up to four days and IL-16 levels were determined over time by ELISA (Fig. 5K). All three sweeping antibodies significantly reduced IL-16 levels compared to WT/Fc^WT^ at both timepoints (Fig. 5L). At the later timepoint, WT/Fc^SW^ showed similar degradation to both L:Y53/Fc^SW^ and H:Y73/Fc^SW^, consistent with its increased recycling capacity compared to WT/Fc^WT^. However, at both early and late timepoints, H:G50/Fc^SW^ demonstrated superior IL-16 degradation compared to all other constructs.

Taken together, we demonstrate the successful generation of sweeping anti-IL-16 antibodies, validate their binding to hIL-16 and FcRn, demonstrate FcRn-mediated cell binding and recycling, and demonstrate efficient short-term and long-term degradation of IL-16 by 14.1-H:G50.

## DISCUSSION

In recent years, there has been an increase in the number of monoclonal antibodies targeting soluble ligands that have been FDA-approved or are currently in development^35^. Soluble ligands are an attractive class of therapeutic targets because of their accessibility^36^ and easier targetability compared to common but difficult-to-target cell surface receptors^37^. In addition, targeting a single soluble ligand can in some cases alter signaling through multiple receptors and thereby increase therapeutic activity^38,39^. To enhance targeting of soluble ligands, sweeping antibody technology has been developed. Sweeping antibodies contain two modifications to enhance clearance of soluble ligands: (1) enhanced binding of Fc to the neonatal Fc receptor (FcRn), and (2) pH-dependent binding of IL-16. While the number of sweeping antibodies currently in use is relatively small, due to their novelty and relatively complex engineering, the first sweeping antibody, ravulizumab, has recently been approved by the FDA for treatment of diseases associated with increased complement activation^40^.

Here, we sought to develop the first sweeping antibody targeting interleukin 16 (IL-16), a soluble ligand that has been suggested as a new target for the treatment of inflammatory diseases and cancer^17,20,41,42^. Pre-clinically, IL-16 has been successfully targeted in models of acute kidney disease^24^, experimental autoimmune encephalitis^43^, type 1 diabetes^44^, and in combination with colorectal cancer therapies^45^. However, IL-16 neutralization as a therapeutic strategy has not been advanced beyond the pre-clinical stage to date. Due to its high level of expression in disease^27^, achieving clinical efficacy will likely rely on highly efficient IL-16 clearance.

To enable efficient antibody-mediated clearance of IL-16, we attempted to engineer sweeping anti-IL-16 antibodies. Using an empirical mutagenesis strategy focused on mutating complementarity-determining region (CDR) amino acid residues to histidine, we were able to successfully introduce pH-dependent binding, creating an acid switch, into the anti-IL-16 antibody clone 14.1 but not clone 3B6. Through epitope analysis of the two antibodies, we identified key differences in the epitope that may confer amenability to the introduction of pH-dependent binding. In addition, we show that in the presence of FcRn-expressing cells, 14.1-based sweeping antibodies more efficiently cleared IL-16 than the parental non-sweeping antibody.

Different methods have been employed to generate pH-dependent antibodies^46,47^. Here, we used a method focused on engineering parental antibodies by introducing single-, double- or multi-histidine mutations into the CDRs. Three single-histidine mutants based on 14.1 exhibited potent acid switch activity while 3B6 remained largely pH-independent despite multiple histidine substitutions. Consequently, we deemed 3B6 resistant to pH-dependent mutations. However, this finding could be the result of limiting the search space to the CDRs. Other approaches, such as predictive algorithms^48^ to determine the best site for histidine replacement across the full variable region (CDRs and frameworks), could prove more successful.

As part of this work, we established a robust workflow for the evaluation of pH-dependent antibody binding by bio-layer interferometry. The key factors that we identified for the successful performance of this assay were: use of hIL-16 as ligand, antibodies in single-arm Fab format as analyte, and all steps of the assay occurring in buffer adjusted to the respective pH. Other assay setups and orientations may be possible and even necessary for other molecules, but the described experimental workflow was crucial to yield robust data in our hands.

To better understand the potential factors that could have led to 14.1 but not 3B6 being amenable to pH-dependent binding, we decided to first determine the 3B6 epitope in addition to the known epitope of 14.1. In this analysis, we discovered that the two antibodies bound to the same general region of hIL-16 leading to partial competition. This presented us with a unique opportunity to assess factors of the epitope that determine amenability to the introduction of pH-sensitivity. We assessed the area of the epitope, its electrostatic potential, as well as structural differences at the site of mutation for the most pH-dependent 14.1-based sweeping antibody, H:G50. For most of the comparisons, we relied on deletion mutants for experimental epitope mapping as well as predictive algorithms to establish epitope characteristics, including epitope size and electrostatic potential. We hypothesize that the smaller epitope area as well as proximity of the His^50^ side chain to a positively charged residue on IL-16, Arg^86^, contributed to the potent acid switch mechanism in this clone. For 3B6, we hypothesize that the larger epitope area of 3B6 provides redundancy through additional contact residues resulting in its relative resistance to creating a molecular switch. While predictive algorithms have become highly accurate in modeling molecular structures^49,50^, experimental structural analysis using NMR and/or isothermal titration calorimetry^51^ of the antibodies alone or in complex with IL-16 at different pH levels may allow for confirmation of the predicted structural changes, inter- and intra-molecular bond formation and protonation states of side chains that would lead to pH-dependent binding. Deepening our understanding of the properties enabling the development of this growing class of therapeutic molecules will be crucial for future antibody development.

In conclusion, in this study, we set out to generate the first sweeping antibodies against IL-16 by engineering conventional antibodies into sweeping antibodies to more efficiently clear IL-16. We were able to identify several parameters correlated with amenability to the introduction of pH-dependent binding and created a sweeping antibody that demonstrated significantly increased clearance of IL-16 in vitro. The successful targeting of cancer- and autoimmunity-associated cytokines using monoclonal antibodies likely requires highly efficient degradation to confer therapeutic activity and our work provides a critical step towards establishing such an approach targeting IL-16.

## MATERIALS AND METHODS

### Cell lines and primary human cells

HMEC-1 cells were purchased from American Type Culture Collection (ATCC), Expi293 cells were purchased from Thermo, and Lenti-X 293T cells were purchased from Takara. Cell lines were cultured according to the manufacturer’s instructions. Cell lines were authenticated by their respective suppliers.

Competent TG1 cells, BL21(λDE3)pBirA cells, and Stbl3 cells were obtained from Lucigen, AMID biosciences, and ThermoFisher (respectively).

Healthy donor buffy coats were obtained from the Blood Centers of America and the New York Blood Center. Healthy donor peripheral blood mononuclear cells (PBMC) were isolated from healthy donor buffy coats via density gradient using FicollPaque (GE) as previously described^52^.

### Expression and isolation of hIL-16, hIL-16 deletion mutants, Fabs and antibodies in Expi293 cells

Expi293 cells were cultured and transfected using the ExpiFectamine™ 293 Transfection kit (ThermoFisher, catalog no. A14524) according to manufacturer’s protocols. To generate IgGs and Fabs, we transfected cells with heavy and light chain-encoding plasmids (pcDNA3.4 backbone) at a 2:1 ratio. To express biotinylated human IL-16 (hIL-16), we transfected cells with an hIL-16 encoding plasmid (pcDNA3.4 backbone) and the secreted-BirA-FLAG plasmid (Addgene plasmid no. 64395) at a 2:1 ratio. Cells were incubated overnight at 37°C, 8% CO_2_ at 125 rpm for 25 ml cultures or 230 rpm for 2.5 ml cultures. If proteins were biotinylated, 50 mM biotin was added to cultures. One day after transfection, ExpiFectamine enhancers were added to each culture. All proteins were harvested on day 6. hIL-16 (PRO_0000015412) and Fabs were isolated using HisPur™ Ni-NTA resin (ThermoFisher, catalog no. 88221). Fc-containing antibodies were isolated using Protein G agarose (Invitrogen, catalog no. 15920-010). All proteins were dialyzed against 1x PBS overnight at 4°C with the Pur-A-Lyzer Dialysis kits (SigmaAldrich, catalog no. PURX35015/PURN60030).

### Single-chain variable fragment (scFv) phage selection

A panning selection was performed using a previously constructed human antibody phage display library^52^ on recombinant human IL-16 followed by recombinant mouse IL-16 (Sino Biological, catalog no. 51303-M07E). Both antigens were immobilized at 10 μg/ml on MaxiSorp Immunotubes (Nunc). Enrichment of anti-human-IL-16-specific phage binders was confirmed using a time-resolved fluorescence immunoassay (TRFIA). For light chain shuffling, selected V_H_ regions were cloned into a diverse light-chain phagemid library. The resulting light-chain shuffled library was used in two subsequent rounds of solution-phased selections using limiting antigen concentrations (10 nM, 1 nM or 100 pM) of hIL-16.

### ELISA – Binding of 14.1 to hIL-16 and mIL-16

High binding ELISA plates (Corning, catalog no. 3361) were coated with 50 μl/well of 10 <u>m</u>g/ml parental 14.1 antibody (Biolegend, catalog no. 519108) in PBS. The plate was incubated overnight at 4°C and subsequently blocked with 4% BSA/PBS. Biotinylated hIL-16 or mIL-16 were added to wells in technical triplicates with a highest concentration of 32.8 μg/ml and 2x dilutions. Plates were incubated for 1 hour at RT. IL-16 binding was detected via 1:1,000 dilution of streptavidin-HRP (Biolegend, catalog no. 405210). TMB was used as the chromogenic substrate. Reaction was stopped with 1 M H_2_SO_4_. Absorbance was read at 450 nm with a reference wavelength at 570 nm using a Spark multi-mode plate reader (Tecan).

### Expression of scFvs and hIL-16 binding

ScFv sequences were cloned into the bacterial expression plasmid, pSANG10-FLAG-Avitag. scFvs were expressed in BL21(λDE3)pBirA cells overnight in Magic Media™ (ThermoFisher) with 50 mM biotin at 37°C, 225 rpm. Supernatants were allowed to bind to immobilized recombinant hIL-16 for 1 h and bound scFvs were detected using an anti-FLAG antibody (SigmaAldrich) followed by an Eu-N1-anti-mouse IgG (PerkinElmer). Fluorescence was read after addition of DELFIA™ Enhancement solution (PerkinElmer) using a Spark multi-mode plate reader (Tecan).

### Conversion of binders to single-arm Fabs

Heavy and light chain variable regions were PCR-amplified and splice-by-overlap-extension PCR was used to join light and heavy variable domains with their respective constant chains and chains were cloned into pcDNA3.4. The heavy chain constant region contained a hexahistidine tag for purification for the Fabs.

<u>Heavy chain IgG1 CH1:</u>

TKGPSVFPLAPSSKSTSGGTAALGCLVKDYFPEPVTVSWNSGALTSGVHTFPAVLQSSGLYSLSSVVTVPSSSLGTQTYICNVNHKPSNTKVDKKVEPHHHHHH*

<u>Kappa light chain constant:</u>

RTVAAPSVFIFPPSDEQLKSGTASVVCLLNNFYPREAKVQWKVDNALQSGNSQESVTEQDSKDSTYSLSSTLTLSKADYEKHKVYACEVTHQGLSSPVTKSFNRGEC*

<u>Lamba light chain constant:</u>

GQPKANPTVTLFPPSSEELQANKATLVCLISDFYPGAVTVAWKADGSPVKAGVETTKPSKQSNNKYAASSYLSLTPEQWKSHRSYSCQVTHEGSTVEKTVAPTECS*

Whole Plasmid Sequencing of plasmids was performed by Plasmidsaurus using Oxford Nanopore Technology with custom analysis and annotation.

### Bio-layer interferometry (BLI) at pH 7 and at pH < 6

Streptavidin (SA) biosensors (Sartorius, catalog no. 18-5020) were prepared in 1x Octet® kinetics buffer (Sartorius, catalog no. 18-1105) for 10 minutes at room temperature prior to use. Loading of sensors and kinetic runs were performed on the Octet K2 (Sartorius). SA biosensors were dipped into baseline wells of 1x Octet® kinetics buffer for 1 minute prior to loading of ligands. Biotinylated ligands, prepared in 1x Octet® kinetics buffer, were loaded onto the SA biosensors for 120-180 seconds at the indicated concentrations (Suppl. Table 1) based on prior ligand scouting experiments, per Sartorius protocols. Following loading, SA biosensors were immediately blocked with 10 μM biocytin in 1x Octet® kinetics buffer for 30 seconds. For the association step, loaded SA biosensors were moved into wells of the analyte. Serial dilutions of analytes were made in 1x Octet® kinetics buffer (Suppl. Table 1). The association step ranged from 120-180 seconds. Immediately following association, SA biosensors were moved into dissociation wells containing 1x Octet® kinetics buffer for 180-600 seconds. For BLI at pH < 6 all steps were carried out as previously described but with 1x Octet® kinetics buffer adjusted to pH < 6 using HCl. BLI was run at 30°C and 1,000 rpm. Data were analyzed using Octet® K2 System Analysis 9.0 Software. Criteria for analyzing binding curves was for χ^2^ ≈ 3 and R^2^ ^>^ 0.95.

### ELISA – Binding of 3B6 to HEL or IL-16

A high binding ELISA plate (Corning, catalog no. 3361) was coated with 50 μl/well of 10 μg/ml hen egg lysozyme (HEL, Rockland, catalog no. MB-109-1000) or hIL-16. Plate was incubated overnight at 4°C. Plate was then blocked with 4% BSA/PBS. 3B6 Fab was added to each well in technical triplicate with a top concentration of 25 μg/ml and a 2-fold dilution series. Plate was incubated for 1 hour at RT. 3B6 binding was detected with 1:5,000-diluted peroxidase-conjugated anti-human IgG F(ab’)2 (Jackson Laboratory, catalog no. 109-035-097). TMB was used as the chromogenic substrate. Reaction was stopped with 1 M H_2_SO_4_. Absorbance was read at 450 nm with a reference wavelength at 570 nm using a Spark multi-mode plate reader (Tecan).

### ELISA – Titration binding curve of 14.1 IgG to hIL-16 and mIL-16

A high binding ELISA plate (Corning, catalog no. 3361) was coated with 50 μl/well 10 μg/ml hIL-16 or mIL-16. Plate was incubated overnight at 4°C. Plate was then blocked with 4% BSA/PBS. Biotinylated hIL-16 was prepared in 4% BSA/PBS at 32.8 mg/ml and a 2-fold dilution series. Biotinylated hIL-16 binding was detected with 1:1000-diluted streptavidin-HRP (Biolegend, catalog no. 405210). TMB was used as the chromogenic substrate. Reaction was stopped with 1 M H_2_SO_4_. Absorbance was read at 450 nm with a reference wavelength at 570 nm using a Spark multi-mode plate reader (Tecan).

### Flow cytometry

Donor PBMCs were thawed one day prior to use. PBMCs were activated using 5 ng/ml PMA and 500 ng/ml ionomycin and incubated at 37°C/5%CO_2_ for 3 hours. Activated PBMCS were washed and incubated with AF488-labeled (ThermoFisher, cat. no. A30006) IL-16, IL-16/AF488 pre-incubated for 1 hour at RT with 14.1 IgG (molar ratio of 1:1), or IL-16/AF488 pre-incubated for 1 hour at RT with 3B6 Fab (molar ratio of 1:1) for 30 minutes at 37°C/5%CO_2_. PBMCs were washed and stained with anti-CD3/BV650 and anti-CD69/PE-Cy7 and incubated at 4°C for 30 minutes. After incubation, cells were washed and ran on a LSR II flow cytometer (BD). Data were analyzed using FlowJo 10. Antibodies used are listed in Suppl. Table 2.

### Protein concentration quantification and SDS-PAGE

Protein concentrations were determined using the Pierce™ BCA protein Assay Kit (ThermoFisher, catalog no. 23227). To assess purity by gel electrophoresis, all proteins were run under reducing, denaturing conditions, except for 14.1 1xH IgGs. 14.1 1xH IgGs were run under non-reducing conditions. All proteins were run on Bolt™ 4-12%, Bis-Tris Plus WedgeWell™ Gels (Invitrogen, catalog no. NW04125BOX) at 150 V for 1 hour. Gels were stained with protein staining reagent (Bulldog Bio, catalog no. AS001000) for 1 hour and destained in distilled water overnight. Gels were visualized with the iBright 1500 imaging system (ThermoFisher).

### ELISA – 14.1 and 3B6 1xH variants binding to hIL-16 at neutral pH

A high binding ELISA plate (Corning, catalog no. 3361) was coated with 50 μl/well 10 μg/ml hIL-16. Plate was incubated overnight at 4°C. Plate was then blocked with 4% BSA/PBS. 14.1 1xH IgG and 3B6 1xH Fabs were prepared at 25 μg/ml in 4% BSA/PBS and added to the plate in technical duplicate. Antibody and Fab binding were detected with 1:5,000-diluted peroxidase-conjugated anti-Human IgG F(ab’)2 (Jackson Laboratory, catalog no. 109-035-097). TMB was used as the chromogenic substrate. Reaction was stopped with 1 M H_2_SO_4_. Absorbance was read at 450 nm with a reference wavelength at 570 nm using a Spark multi-mode plate reader (Tecan).

### Time-resolved fluorescence immuno-assay – 14.1 1xH variants binding to hIL-16 at neutral pH and pH < 6

A high binding black flat-bottom plate (Greiner, catalog no. 655077) was coated with 50 μl/well 5 μg/ml hIL-16. Plate was incubated overnight at 4°C. Plate was then blocked with 4% BSA/PBS. Biotinylated 14.1 1xH IgG were prepared at 10 μg/ml in 4% BSA/PBS pH 7.4 or 4% BSA/PBS pH < 6 (adjusted using 1 M HCl). 14.1 1xH IgG binding was detected using 1:1,000-diluted streptavidin-Eu (PerkinElmer, catalog no. 1244-360). Time-resolved fluorescence was read after addition of DELFIA™ Enhancement solution (PerkinElmer) using a Spark multi-mode plate reader (Tecan).

### ELISA – 3B6 1xH, 2xH and nxH variants binding to hIL-16 at neutral pH and pH < 6

A high binding ELISA plate (Corning, catalog no. 3361) was coated with 50 μl/well 10 μg/ml hIL-16. Plate was incubated overnight at 4°C. Plate was then blocked with 4% BSA/PBS. 3B6 1xH Fabs were prepared at 25 μg/ml in 4% BSA/PBS pH 7.4 or 4% BSA/PBS pH < 6 (adjusted using 1 M HCl). Fab binding was detected with 1:5,000-diluted peroxidase-conjugated anti-human IgG F(ab’)2 (Jackson Laboratory, catalog no. 109-035-097). TMB was used as the chromogenic substrate. Reaction was stopped with 1 M H_2_SO_4_. Absorbance was read at 450 nm with a reference wavelength at 570 nm using a Spark multi-mode plate reader (Tecan).

### Tandem BLI assay

Streptavidin (SA) biosensors (Sartorius, catalog no. 18-5020) were prepared in 1x Octet® kinetics buffer (Sartorius, catalog no. 18-1105) for 10 minutes at room temperature prior to use. Loading of sensors and kinetic runs were performed on the Octet K2 (Sartorius). SA biosensors were dipped into baseline wells of 1x Octet® kinetics buffer for 30 seconds prior to loading of ligands. Biotinylated rhIL-16, prepared in 1x Octet® kinetics buffer, was loaded onto the SA biosensors for 300 seconds at 1.25 μg/ml. Following loading, rhIL-16-coated sensors were dipped into 1x Octet® kinetics buffer for 30 seconds. Following this baseline step, both sensors were dipped into 250 nM 3B6 Fab for 600 seconds. Immediately following 3B6 Fab loading, one sensor was dipped into a well of 250 nM 3B6 Fab or a well of pre-mixed 250 nM 3B6 Fab and 250 nM 14.1 Fab prepared in 1x Octet® kinetics buffer for 300 seconds. BLI was run at 30°C and 1,000 rpm. Data were analyzed using Octet® K2 System Analysis 9.0 Software.

### Linear epitope scanning for 3B6 IgG against hIL-16

The 121 amino acid sequence of hIL-16 was printed in 15 amino acid peptides with 14 amino acids overlapping using PEPperMAP® Linear Epitope Mapping Microarray (PEPperPRINT® GmbH, Heidelberg, Germany). An IL-16 peptide microarray was pre-stained with the secondary antibody to determine possible background interactions with the linear peptides of the microarray that could interfere with the main assays. Simultaneous incubation of further IL-16 peptide microarrays with human monoclonal IgG 3B6 sample at concentrations of 1 μg/ml, 10 μg/ml and 100 μg/ml was followed by staining with the secondary antibody and read-out with an Innopsys InnoScan 710-IR Microarray Scanner. The additional HA peptides framing the IL-16 peptide microarrays were simultaneously stained with the control antibody as internal quality control to confirm the assay performance and the peptide microarray integrity. Microarray image analysis was done with PepSlide^®^ Analyzer. Intensity maps were generated. A maximum spot-to-spot deviation of 40 was tolerated, otherwise the corresponding intensity value was zeroed.

### AlphaFold3 predictions of 3B6 Fab, 14.1 H:G50 Fab, and hIL-16

For prediction of the 3B6 Fab, the heavy and light chains were entered as separate entities in the AlphaFold3 server. For prediction of hIL-16 structure, the 121 amino acid sequence was entered into the AlphaFold3 server. For both structures, the best prediction based off pTM value was used for visualization and evaluation in ChimeraX as well as docking predictions. For analysis of 14.1 and 14.1 H:G50 in complex with hIL-16 interface, the AlphaFold3-generated 14.1 H:G50 structure was superimposed onto the existing structure (PDB:5FB8) - which features a truncated version of hIL-16 - using the “matchmaker’ command in ChimeraX.

### HADDOCK docking of 3B6 Fab with hIL-16

The PDB files of AlphaFold3 predicted proteins were submitted to the HADDOCKv2.4 server as “Protein or Protein-Ligand” molecules for docking. For 3B6 Fab/hIL-16, the 3B6 CDRs were defined as active residues and all hIL-16 residues were defined as active. Standard docking parameters for the EASY access level were used. For analysis of 14.1 and hIL-16 interface, the AlphaFold3-generated hIL-16 structure was superimposed onto the existing structure (PDB:5FB8) using the “matchmaker’ command in ChimeraX.

### Epitope area and distance measurements

AlphaFold3 generated hIL-16 was visualized in ChimeraX. Epitope residues were highlighted. A new surface containing the three most distant atoms from the central hydrophobic pocket was created using the *shape triangle atoms* command. The area of the newly created surfaces was calculated using the *measure area* command.

AlphaFold3 predicted 14.1 H:G50 Fab was aligned to PDB: 5FB8 (14.1:trhIL-16) in ChimeraX. The atoms His^50^ ND1 of 14.1 H:G50 and Arg^86^ NH1of trhIL-16 were selected and the *distance* tool was used to calculate the distance between the atoms.

### APBA-PDB2PQR analysis of hIL-16 electrostatic potential

The PDB file of AlphaFold-generated hIL-16 was uploaded to the Adaptive Poisson-Boltzmann Solver (APBA)-PDB2PQR software suite^50^ and the continuum electrostatic calculations were determined at pH 7 and pH 5. PyMol structures were downloaded, converted into PDB files and visualized using ChimeraX.

### Conversion into sweeping antibodies

Sweeping antibody constructs were synthesized by Twist Bioscience. Single-arm Fab constructs were converted into full IgG constructs using the following Fc regions.

<u>Fc^WT^ domain:</u>

KSCDKTHTCPPCPAPELLGGPSVFLFPPKPKDTLMISRTPEVTCVVVDVSHEDPEVKFNWYVDGVEVHNAKTKPREEQYNSTYRVVSVLTVLHQDWLNGKEYKCKVSNKALPAPIEKTISKAKGQPREPQVYTLPPSREEMTKNQVSLTCLVKGFYPSDIAVEWESNGQPENNYKTTPPVLDSDGSFFLYSKLTVDKSRWQQGNVFSCSVMHEALHNHYTQKSLSLSPG*

<u>Fc^SW^ domain:</u>

KSCDKTHTCPPCPAPELLGGPSVFLFPPKPKDTLYISRTPEVTCVVVDVSHEDPEVKFNWYVDGVEVHEAKTKPREEQYNSTYRVVSVLTVLHQDWLNGKEYKCKVSNKALPAPIEKTISKAKGQPREPQVYTLPPSREEMTKNQVSLTCLVKGFYPSDIAVEWESNGQPENNYKTTPPVLDSDGSFFLYSKLTVDKSRWQQGNVFSCSVMHEALHYHYTQKSLSLSPG*

### ELISA – Binding of 14.1 sweeping IgGs to hIL-16 and mIL-16

A high binding ELISA plate (Corning, catalog no. 3361) was coated with 50 μl of 10 μg/ml of hIL-16 or mIL-16. Plate was incubated overnight at 4°C. Plate was then blocked with 4% BSA/PBS. Antibodies were prepared in 4% BSA/PBS at 5 μg/ml. Antibodies were detected with 1:1,000-diluted peroxidase-conjugated anti-human IgG-FcY antibody (Jackson Laboratory, catalog no. 109-036-170). TMB was used as the chromogenic substrate. Reaction was stopped with 1 M H_2_SO_4_. Absorbance was read at 450 nm with a reference wavelength at 570 nm using a Spark multi-mode plate reader (Tecan).

### Lentiviral transduction and validation of HMEC-1 cells

HMEC-1 cells were transduced with pTwist-Lenti-SFFV-FcRnα-P2A-β2M-T2A-BFP and sorted for BFP expression on a FACSAria II cell sorter (BD). 7.5 x 10^5^ cells were lysed in RIPA buffer (ThermoFisher, catalog no. 89901) containing protease inhibitor (Roche, catalog no. 04693159001). Protein concentration was determined via Pierce™ BCA protein Assay Kit (ThermoFisher, catalog no. 23227). 30 μg of cell lysates was separated by sodium dodecyl sulfate-gel electrophoresis at 150 V for 40 minutes. Proteins were then transferred to nitrocellulose membranes using an iBlot2 transfer system (ThermoFisher). The membrane was blocked with 5% nonfat milk-tris-buffered saline and incubated with primary antibodies against FcRnα, β2M, and β-actin. Membranes were washed and developed with an anti-mouse-IgG/Horseradish peroxidase antibody and Western Lightning Plus-ECL solution (PerkinElmer). Bands were visualized with the iBright 1500 imaging system (ThermoFisher).

### In vitro antibody recycling assay

The sweeping antibody recycling assay was performed largely as previously described^34^. HMEC-1-BFP^+^ cells were plated in 24-well plates at 1 x 10^5^ cells/well in complete media and incubated at 37°C, 5% CO_2_ overnight. Fresh media was added to each well on the next day and plate was again incubated overnight at 37°C, 5% CO_2_.

For the “Wash” condition, cells were then washed with PBS and incubated in Hank’s Balanced Salt Solution (HBSS, ThermoFisher, catalog no. J67799.K2) for one hour at 37°C, 5% CO_2_. 5 nM antibodies and 5 nM biotinylated IL-16 were added to each well. Plate was incubated at 37°C, 5% CO_2_ for four hours. Cells were then washed four times with ice cold PBS. Cells were then incubated in complete media without FCS overnight. Supernatants were collected the next day.

For the “No wash” condition, antibodies and biotinylated IL-16 were prepared in complete HMEC-1 media and added to each well. Cells were incubated at 37°C, 5% CO_2._ The next day, cell plates were centrifuged at 400 x g for 5 minutes and aliquots of supernatants were collected.

To quantify antibodies in the supernatants, an antibody ELISA was performed. A high binding ELISA plate (Corning, catalog no. 3361) was coated with 50 μl of 10 μg/ml AffiniPure™ F(ab’)_2_ Fragment Donkey Anti-Human IgG (H+L) min. cross react. (Jackson Laboratory, catalog no. 709-006-149). The plate was incubated at 4°C overnight and then blocked with 4% BSA/PBS. Supernatants were added to the wells and antibodies detected with 1:5,000 peroxidase-conjugated anti-human IgG (Jackson Laboratory, catalog no. 109-036-170). TMB was used as the chromogenic substrate. Reaction was stopped with 1 M H_2_SO_4_. Absorbance was read at 450 nm with a reference wavelength at 570 nm using a Spark multi-mode plate reader (Tecan).

### In vitro IL-16 degradation assay

HMEC-1-BFP^+^ cells were plated in 24-well plates at 1 x 10^5^ cells/well in complete media and incubated at 37°C, 5% CO_2_ overnight. 5 nM antibodies and 5 nM biotinylated IL-16 were prepared in complete HMEC-1 media and added to each well. Cells were incubated at 37°C, 5% CO_2._ Each subsequent day, cell plates were centrifuged at 400 x g for 5 minutes, aliquots of supernatants were collected, and remaining supernatants were moved to fresh HMEC-1-BFP^+^ cells plated the day prior. An IL-16 ELISA was performed to detect biotinylated IL-16 in supernatants. A high binding ELISA plate (Corning, catalog no. 3361) was coated with 50 μl/well of 8 μg/ml of anti-IL-16 capture antibody (R&D, catalog no. MAB316-100). The plate was incubated at 4°C overnight. Plate was blocked with 4% BSA/PBS and supernatants were added to the wells. Biotinylated IL-16 was detected via 1:1,000 streptavidin-HRP (Biolegend, catalog no. 405210). TMB was used as the chromogenic substrate. Reaction was stopped with 1 M H_2_SO_4_. Absorbance was read at 450 nm with a reference wavelength at 570 nm using a Spark multi-mode plate reader (Tecan).

### Antibodies

All antibodies used can be found in Suppl. Table 2.

### Statistics

Significance of differences in percentage IL-16 binding to T cells and IL-16 binding to 14.1 antibody in the presence of 3B6 or 14.1 were determined by ordinary one-way ANOVA followed by Dunnett’s multiple comparison test. Significance of difference in relative IgG abundance was determined by an ordinary one-way ANOVA followed by Tukey’s multiple comparisons test. Significance of differences in quantification of IL-16 in degradation assay and binding of polyclonal 2^nd^ round-selected scFvs to IL-16 were determined by two-way ANOVA followed by Šídák’s multiple comparisons test. All statistical analyses were performed in Prism 10 (GraphPad Software) and results were considered significant when p ≤ 0.05.

## ACKNOWLEDGEMENTS

This publication was supported by funds through the Maryland Department of Health’s Cigarette Restitution Fund Program (CH-649-CRF) and supported by an NIAID-funded predoctoral fellowship to JMB (T32 AI095190). The authors would like to acknowledge Dr. Jonathan F. Fay for his intellectual contributions to this publication. We thank the UMGCC Flow Cytometry core facility, UMB Translational Genomics Laboratory and Dr. Oleksandr Galkin from Sartorius. The plasmid pSANG10-3F (Addgene plasmid #39264) was a gift from Dr. J. McCafferty. The secreted BirA-Flag plasmid was a gift from Gavin Wright (Addgene plasmid #64395). For HADDOCK: The FP7 WeNMR (project# 261572), H2020 West-Life (project# 675858), the EOSC-hub (project# 777536) and the EGI-ACE (project# 101017567) European e-Infrastructure projects are acknowledged for the use of their web portals, which make use of the EGI infrastructure with the dedicated support of CESNET-MCC, INFN-LNL-2, NCG-INGRID-PT, TW-NCHC, IFCA-LCG2, UA-BITP, TR-FC1-ULAKBIM, CSTCLOUD-EGI, IN2P3-CPPM, SURFsara and NIKHEF, and the additional support of the national GRID Initiatives of Belgium, France, Italy, Germany, the Netherlands, Poland, Portugal, Spain, UK, Taiwan and the US Open Science Grid.

## AUTHOR CONTRIBUTIONS

JMB and TL conceived the project. JMB performed phage display selections, generated and quantified all antibody, Fab and scFv constructs as well as full hIL-16/mIL-16 constructs and deletion mutants for hIL-16, performed bio-layer interferometry (BLI), ELISAs and TRFIAs to assess binding kinetics, performed flow cytometry binding assays, conducted size-exclusion chromatography, generated prediction structures and docking for hIL-16 and 3B6, performed lentiviral transductions, carried out recycling and degradation assays with corresponding ELISAs and wrote the manuscript. AW performed plasmid isolations and bio-layer interferometry experiments. KAD and DA planned experiments and analyzed data. TL supervised all work related to this project. All authors reviewed and approved the manuscript.

