## Supplementary Material for "A novel sweeping antibody exhibits efficient clearance of the cancer- and autoimmunity-associated cytokine interleukin 16"

*Baker et al., 2025*

**SUPPLEMENTARY FIGURES**

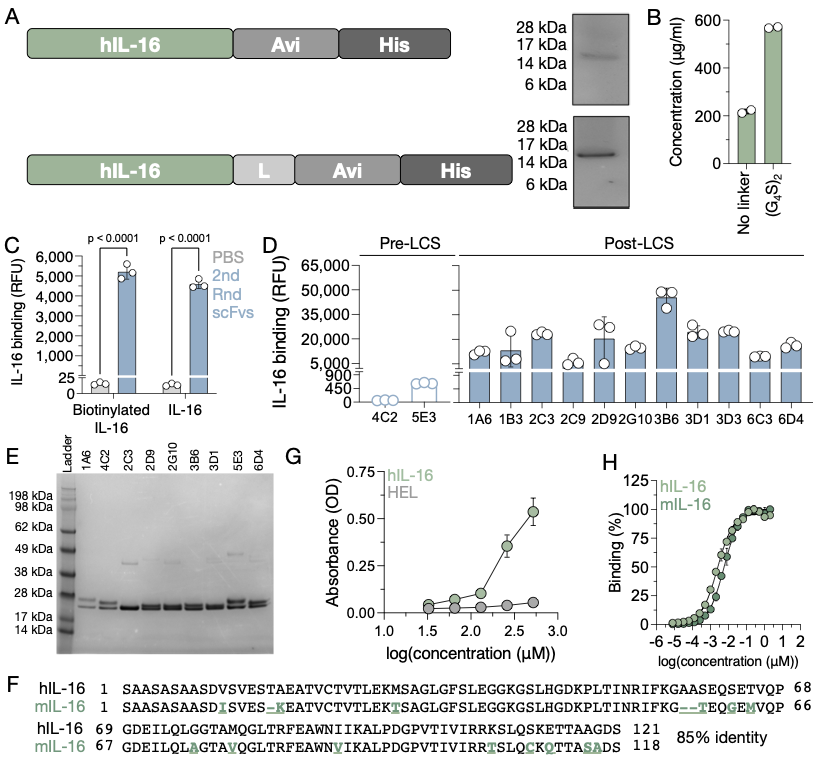
 **Supplementary Figure 1: Expression of IL-16 and binding by anti-IL-16 antibodies. (A)** Schema of human IL-16 (hIL-16) fused to an Avi-tag polypeptide sequence (Avi) and a deca-histidine (His) tag with and without a (Gly_4_Ser)_2_ linker (L). Corresponding SDS-PAGE gel images stained with Coomassie brilliant blue (Bulldog-Bio, cat. no. AS001000) are shown; the predicted molecular weight of hIL-16 with linker is 16.2 kDa and without linker is 15.6 kDa. **(B)** Protein concentrations of both hIL-16 constructs with and without (Gly_4_Ser)_2_ linker following Ni-NTA purification and dialysis into PBS. **(C)** Binding of polyclonal phage derived from round 2 selection outputs hIL-16 with and without biotinylation as determined by time-resolved fluorescence immunoassay (TRFIA). Data indicate means ± SD from technical replicates (n = 3). Statistical differences between the means were determined by a two-way ANOVA followed by Šídák’s multiple comparisons test. **(D)** Binding of two pre-light chain shuffled (LCS) scFvs and eleven post-LCS scFvs from round 4 to human IL-16 (hIL-16) as determined TRFIA. **(E)** Fixed mass (2 ug) of Ni-NTA-purified scFvs visualized on a reducing/denaturing SDS-PAGE gel image. **(F)** BLAST protein sequence alignment of hIL-16 and mouse IL-16 (mIL-16). Differences between hIL-16 and mIL-16 are highlighted in green and underlined. Missing amino acids are represented by a hyphen. **(G)** Binding of 3B6 Fab to hIL-16 of hen egg lysozyme (HEL) as determined by ELISA. Data indicate means ± SD from technical replicates (n = 3). **(H)** Titration curve for binding of 14.1 IgG to hIL-16 and mIL-16. Data indicate means ± SD from technical replicates (n = 2). Data represents binding data from two independent experiments.

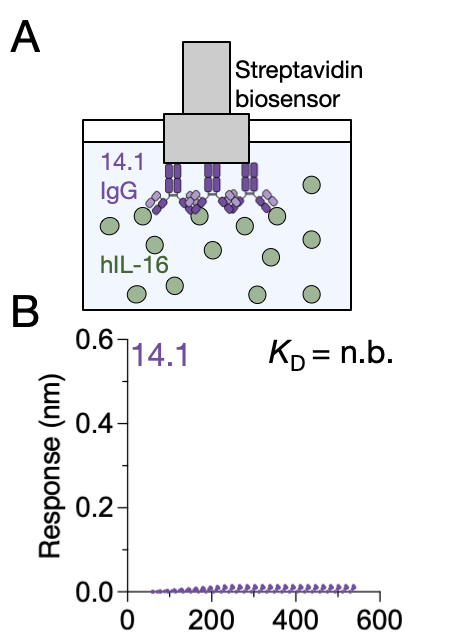

**Supplementary Figure 2: Impact of assay orientation on biolayer interferometry. (A)** Schema of BLI assay set-up for measuring binding kinetics in reverse orientation using IL-16 (hIL-16) as analyte and 14.1 IgG as ligand at an acidic pH (pH < 6). **(B)** BLI sensorgram of 14.1 IgG binding to hIL-16 in reverse assay orientation. Data indicate representative binding from two independent experiments. n.b. = no binding.

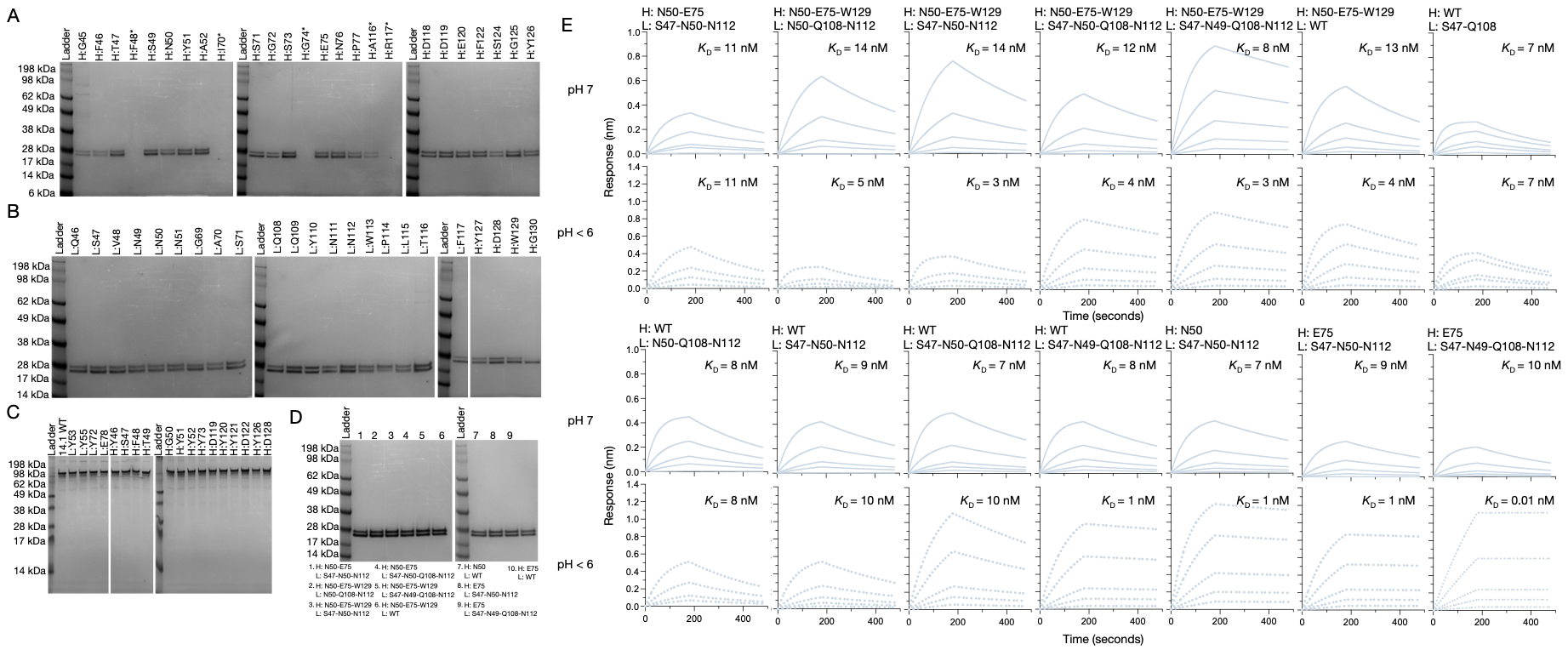

**Supplementary Figure 3: Expression of histidine variant anti-IL-16 antibodies and their binding kinetics.** Reducing and denaturing SDS-PAGE gel of **(A/B)** single-histidine 3B6 Fab heavy and light chain variants loaded at a fixed mass (0.5 μg) or maximum volume (*) following purification via Ni-NTA resin and dialysis into PBS. **(C)** Non-reducing and non-denaturing SDS-PAGE gel of single-histidine 14.1 IgG variants loaded at a fixed mass (2 μg) following purification via Protein G resin and dialysis into PBS. **(D)** Reducing and denaturing SDS-PAGE gel of multi-histidine 3B6 Fab variants loaded at a fixed mass (2 μg) following purification via Ni-NTA resin and dialysis into PBS. **(E)** Bio-layer interferometry (BLI) sensorgrams of 3B6 nxH variant Fabs binding to hIL-16 at neutral (pH 7) and acidic (pH < 6) pH. Data indicate representative result from two independent experiments for each variant.

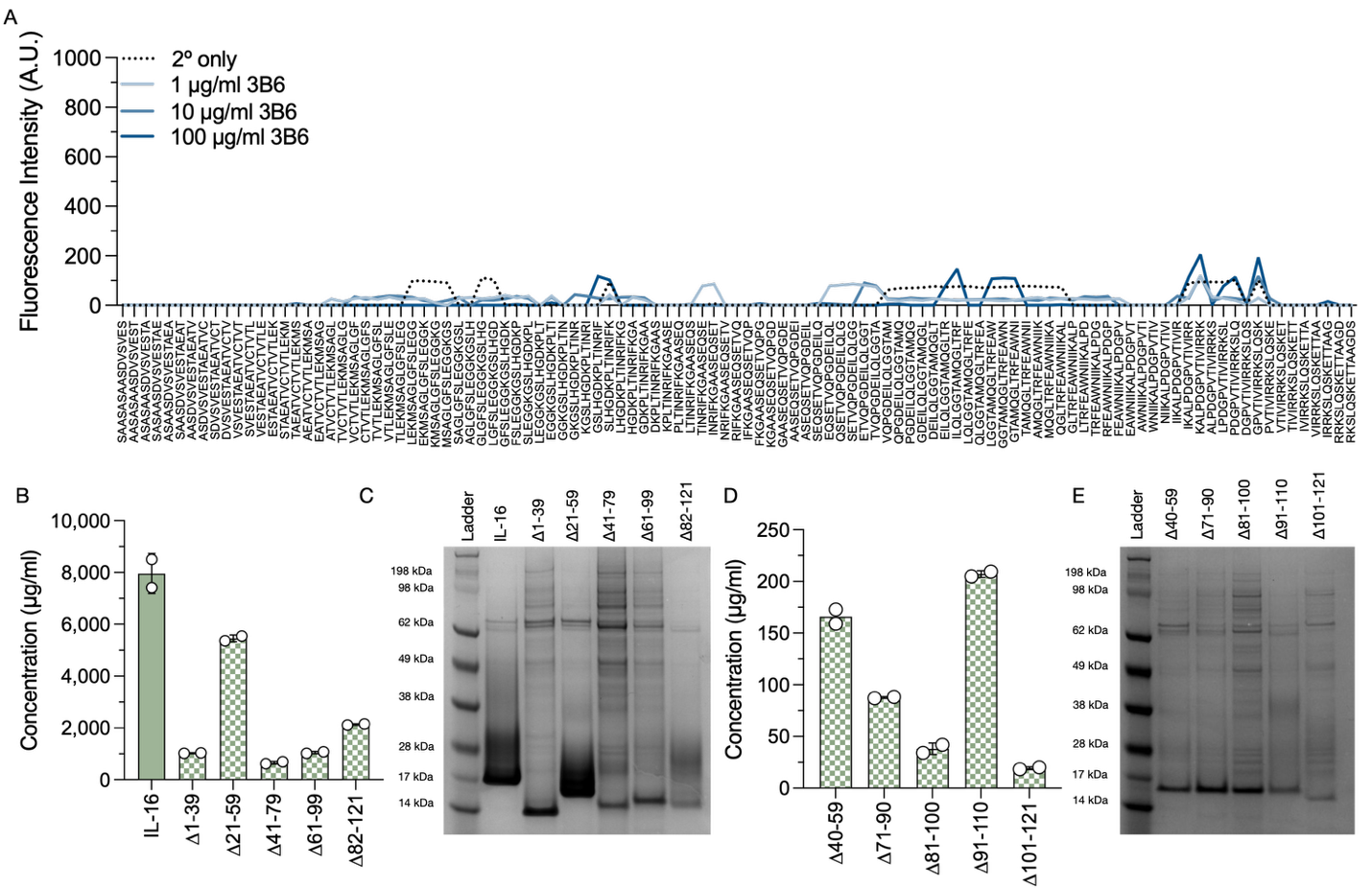

**Supplementary Figure 4: Experimental determination of the 3B6 epitope. (A)** Linear epitope scanning of 3B6 IgG using linear 15-mer overlapping peptides of hIL-16 at three concentrations. Data indicate means from technical replicates (n = 2). **(D)** Protein concentrations and **(E)** reducing and denaturing SDS-PAGE gel of hIL-16 and 40-amino acid deletion mutants of hIL-16. Data indicate means ± SD from technical replicates (n = 2). **(F)** Protein concentration and **(G)** reducing and denaturing SDS-PAGE gel of 20-amino acid deletion mutants of hIL-16. Data indicate means ± SD from technical replicates (n = 2).

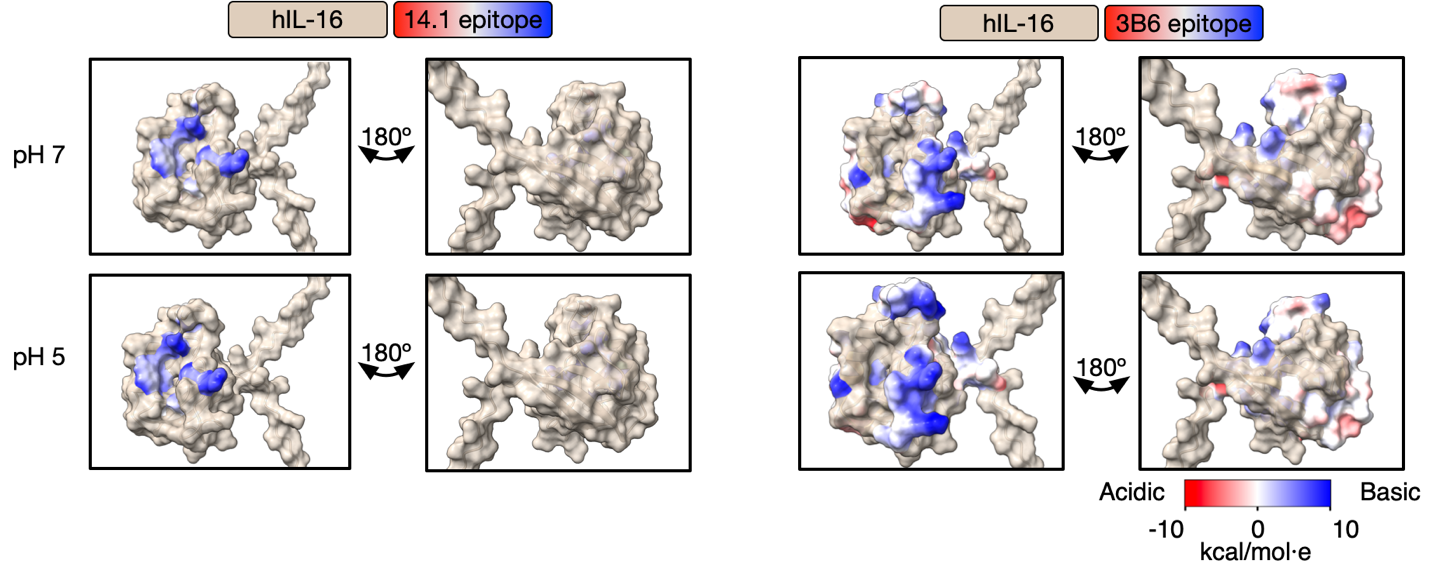
**Supplementary Figure 5: Comparison of the electrostatic potential of anti-IL-16 epitopes.** Electrostatic charge of human IL-16 (hIL-16) (tan) residues that form the 14.1 and 3B6 epitope at both pH 7 and pH 5.

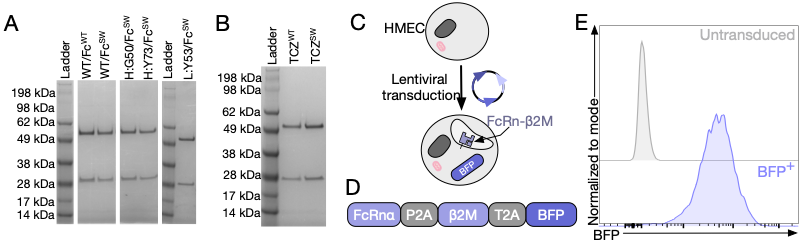

**Supplementary Figure 6: Generation of sweeping antibodies and HMEC-FcRnα-β2M cells. (A)** Reducing and denaturing SDS-PAGE gel of 14.1 sweeping and control antibodies loaded at a fixed mass (1 μg) following purification via Protein G resin and dialysis into PBS. **(B)** Reducing and denaturing SDS-PAGE gel of parental tocilizumab (TCZ^WT^) and sweeping tocilizumab (TCZ^SW^) antibodies loaded at a fixed mass (1 μg) following purification via Protein G resin and dialysis into PBS. **(C)** Schema of lentiviral transduction process and **(D)** lentiviral construct used to generate HMEC-FcRnα-β2M cells. FcRnα = FcRn alpha chain; β2M = β2 microglobulin; BFP = blue fluorescent protein. **(E)** Flow cytometry analysis of BFP expression of untransduced and transduced HMEC cells sorted for BFP expression (BFP^+^).

**SUPPLEMENTARY TABLES**

**Supplementary Table 1.** Parameters for bio-layer interferometry experiments.

| Analyte | Ligand | pH | Ligand Concentration (μg/ml) | Top Analyte Concentration (nM) | Dilution Factor for Analyte |
| --- | --- | --- | --- | --- | --- |
| 4C2 Fab | hIL-16 | 7 | 5 | 2000 | 1.5 |
| 1A6 Fab | hIL-16 | 7 | 5 | 1000 | 1.5 |
| 2G10 Fab | hIL-16 | 7 | 5 | 2000 | 1.5 |
| 3D1 Fab | hIL-16 | 7 | 5 | 2000 | 1.5 |
| 3B6 Fab | hIL-16 | 7 | 1.25 | 111.1 | 1.5 |
| 3B6 Fab | mIL-16 | 7 | 2.5 | 111.1 | 1.5 |
| 14.1 Fab | hIL-16 | 7 | 0.3125 | 111.1 | 1.5 |
| 14.1 Fab | mIL-16 | 7 | 0.3125 | 88.8 | 1.5 |
| 14.1 Fab | hIL-16 | < 6 | 0.3125 | 250 | 1.5 |
| 3B6 Fab | hIL-16 | < 6 | 1.25 | 111.1 | 1.5 |
| hIL-16 | 14.1 IgG | < 6 | 8 | 200 | 2 |
| 3B6 H:N50/L:Q108 Fab | hIL-16 | 7 | 0.625 | 111.1 | 1.5 |
| 3B6 H:N50/L:Q108 Fab | hIL-16 | < 6 | 0.625 | 111.1 | 1.5 |
| 3B6 H:E75/L:S47 Fab | hIL-16 | 7 | 1.25 | 111.1 | 1.5 |
| 3B6 H:E75/L:S47 Fab | hIL-16 | < 6 | 1.25 | 111.1 | 1.5 |
| 3B6 H: N50-E75/L: S47-N50-N112 Fab | hIL-16 | 7 | 1.25 | 62.5 | 2 |
| 3B6 H: N50-E75/L: S47-N50-N112 Fab | hIL-16 | < 6 | 1.25 | 31.3 | 2 |
| 3B6 H: N50-E75-W129/  L: N50-Q108-N112 Fab | hIL-16 | 7 | 1.25 | 62.5 | 2 |
| 3B6 H: N50-E75-W129/  L: N50-Q108-N112 Fab | hIL-16 | < 6 | 1.25 | 31.3 | 2 |
| 3B6 H: N50-E75-W129/L: S47-N50-N112  Fab | hIL-16 | 7 | 1.25 | 62.5 | 2 |
| 3B6 H: N50-E75-W129/L: S47-N50-N112  Fab | hIL-16 | < 6 | 1.25 | 31.3 | 2 |
| 3B6 H: N50-E75-W129/L: S47-N50-Q108-N112 Fab | hIL-16 | 7 | 1.25 | 62.5 | 2 |
| 3B6 H: N50-E75-W129/L: S47-N50-Q108-N112 Fab | hIL-16 | < 6 | 1.25 | 62.5 | 2 |
| 3B6 H: N50-E75-W129  L: S47-N49-Q108-N112 Fab | hIL-16 | 7 | 1.25 | 62.5 | 2 |
| 3B6 H: N50-E75-W129  L: S47-N49-Q108-N112 Fab | hIL-16 | < 6 | 1.25 | 62.5 | 2 |
| 3B6 H: N50-E75-W129/L: WT  Fab | hIL-16 | 7 | 1.25 | 62.5 | 2 |
| 3B6 H: N50-E75-W129/L: WT  Fab | hIL-16 | < 6 | 1.25 | 62.5 | 2 |
| 3B6 H: WT/L: S47-Q108 Fab | hIL-16 | 7 | 1.25 | 62.5 | 2 |
| 3B6 H: WT/L: S47-Q108 Fab | hIL-16 | < 6 | 1.25 | 62.5 | 2 |
| 3B6 H: WT/L: N50-Q108-N112 Fab | hIL-16 | 7 | 1.25 | 62.5 | 2 |
| 3B6 H: WT/L: N50-Q108-N112 Fab | hIL-16 | < 6 | 1.25 | 31.3 | 2 |
| 3B6 H: WT/L: S47-N50-N112 Fab | hIL-16 | 7 | 1.25 | 62.5 | 2 |
| 3B6 H: WT/L: S47-N50-N112 Fab | hIL-16 | < 6 | 1.25 | 62.5 | 2 |
| 3B6 H: WT/L: S47-N50-Q108-N112 Fab | hIL-16 | 7 | 1.25 | 62.5 | 2 |
| 3B6 H: WT/L: S47-N50-Q108-N112 Fab | hIL-16 | < 6 | 1.25 | 62.5 | 2 |
| 3B6 H: WT/L: S47-N49-Q108-N112 Fab | hIL-16 | 7 | 1.25 | 62.5 | 2 |
| 3B6 H: WT/L: S47-N49-Q108-N112 Fab | hIL-16 | < 6 | 1.25 | 62.5 | 2 |
| 3B6 H: N50/L: S47-N50-N112 Fab | hIL-16 | 7 | 1.25 | 62.5 | 2 |
| 3B6 H: N50/L: S47-N50-N112 Fab | hIL-16 | < 6 | 1.25 | 62.5 | 2 |
| 3B6 H: E75/L: S47-N50-N112 Fab | hIL-16 | 7 | 1.25 | 62.5 | 2 |
| 3B6 H: E75/L: S47-N50-N112 Fab | hIL-16 | < 6 | 1.25 | 62.5 | 2 |
| 3B6 H: E75/L: S47-N49-Q108-N112 Fab | hIL-16 | 7 | 1.25 | 62.5 | 2 |
| 3B6 H: E75/L: S47-N49-Q108-N112 Fab | hIL-16 | < 6 | 1.25 | 62.5 | 2 |
| 14.1 L:Y53 | hIL-16 | 7 | 0.3125 | 111.1 nM | 1.5 |
| 14.1 L:Y53 | hIL-16 | < 6 | 0.3125 | 250 nM | 1.5 |
| 14.1 H:G50 | hIL-16 | 7 | 0.3125 | 111.1 nM | 1.5 |
| 14.1 H:G50 | hIL-16 | < 6 | 0.3125 | 250 nM | 1.5 |
| 14.1 H:Y73 | hIL-16 | 7 | 0.3125 | 74.1 nM | 1.5 |
| 14.1 H:Y73 | hIL-16 | < 6 | 0.3125 | 250 nM | 1.5 |

**Supplementary Table 2.** Complete list of antibodies used in experiments.

| **Antibodies** | **Manufacturer** | **Category Number** | **Clone** | **Use** |
| --- | --- | --- | --- | --- |
| Mouse anti-FLAG | Sigma Aldrich | F3165-1MG | M2 | TRFIA |
| DELFIA™ Eu-N1-anti-mouse IgG | PerkinElmer | AD0207 |  | TRFIA |
| Peroxidase-conjugated AffiniPure Goat Anti-Human IgG F(ab’)2 fragment specific | Jackson Laboratory | 109-035-097 | Polyclonal | ELISA |
| Anti-CD3/BV650 | Biolegend | 300468 | UCHT1 | Flow |
| Anti-CD69/PE-cy7 | Biolegend | 310912 | FN50 | Flow |
| Anti-human-FcRn | R&D | MAB8639 | 937508 | Western |
| Anti-human-β2M | R&D | MAB8248 | 883028 | Western |
| Anti-Human/Mouse/Rat-β-actin | R&D | MAB8929 | 937215 | Western |
| Anti-mouse IgG-HRP | R&D | HAF007 | Polyclonal | Western |
| Peroxidase-conjugated AffiniPure F(ab’)2 Goat Anti-Human IgG, FcY Fragment specific | Jackson Laboratory | 109-036-170 | Polyclonal | ELISA, Recycling assay |
| AffiniPure™ F(ab’)_2_ Fragment Donkey Anti-Human IgG (H+L) min. cross react. | Jackson Laboratory | 709-006-149 | Polyclonal | Recycling assay |
| Anti-human-IL-16 antibody | R&D | MAB316-100 | 70719 | Degradation assay |
